# GPCR-Gα13 signaling regulates survival of intestinal intraepithelial CD8 lymphocytes through migration to cytokine-rich niches

**DOI:** 10.64898/2026.03.01.708910

**Authors:** Zachary M. Earley, Anshul Rao, Longhui Qiu, Konrad Knöpper, Fanglue Peng, Norihide Jo, Ananya Krishnapura, Wioletta Lisicka, Hanna Taglinao, Jinping An, Ying Xu, Li V. Yang, Dan Liu, Mark R. Looney, Jason G. Cyster

## Abstract

Intestinal intraepithelial lymphocytes (IEL), including conventional CD8αβ T resident memory (Trm) cells and unconventional CD8αα T cells, promote tissue integrity. Here, we studied the G-protein coupled receptor signals regulating IEL positioning, homeostasis, and function. Deficiency in heterotrimeric G-protein subunit Gα13 or its effector Arhgef1 caused an intestine-specific loss of all types of CD8^+^ TCRαβ and TCRγδ IEL. Gα13-deficient IEL exhibited restricted intraepithelial movement and impaired maturation. Induction of intestinal CD8αβ^+^ Trm upon infection was intact in the absence of Gα13-signaling, but the cells had poor access to the villous niche and defective survival that could be rescued by increasing TGFβ^+^ or interleukin (IL)15. In vivo CRISPR/Cas9 screening identified GPR132 as a Gα13-coupled receptor that regulates CD8αβ^+^ IEL homeostasis and migration to lysophosphatidylcholine. Mice bearing Gα13-deficient T cells suffered more severe colitis and increased colorectal tumor growth. The selective requirement for Gα13-signaling for IEL positioning and survival in the villous niche has implications for therapeutic intervention.

## INTRODUCTION

Intestinal intraepithelial lymphocytes (IELs) play an essential role in maintaining the intestinal barrier and provide a first line of defense against pathogens and intestinal tumors. There are an estimated 20-50 million IEL in the mouse small intestine.^1^ IELs include CD8 and CD4 T cells, with CD8 cells being predominant. CD8 IELs are divided into so-called unconventional T cells that include CD8αα expressing γδT cells and αβT cells that are programmed in the thymus, and conventional CD8αβ αβT cells that include T resident memory (Trm) cells induced during mucosal immune responses.^1^ Both γδT IEL and αβT IEL are motile and survey the epithelium, contacting multiple epithelial cells per hour.^2–4^ The mechanisms responsible for T cell recruitment to the intestine have been extensively studied, with CCR9 playing an important role responding to CCL25 that is selectively expressed by small intestinal epithelium ^5,6^, and GPR15 responding to GPR15L in the large intestine.^7–9^ The integrin α4β7 functions in T cell access to the small and large intestine and αEβ7 (CD103) contributes to retention in the intestine.^1,5,6,10^ However, the full constellation of factors determining T cell migration within and positioning along the villi are unclear.

While the different IEL subsets occupy a similar niche, they have both shared and distinct requirements for their maintenance. γδT IEL are strongly dependent and CD8αα IEL partially dependent on IL15 and IL15Rα.^1^ CD8αβ IEL are less dependent on this cytokine receptor system alone, though they are reduced in number when both IL15 and IL7 are deficient.^11^ The process of conventional CD8αβ T cells becoming intestinal IEL involves priming in mesenteric lymph nodes (MLN) where retinoic acid (RA) promotes α4β7 and CCR9 expression.^5,12–15^ Following arrival in the intestine, the cells are exposed to locally activated TGFβ leading to a TGFβ-induced gene expression program that includes CD103.^16–19^ Persistence of CD8αβ cells in the intestine beyond a few days requires TGFβR signaling. In the LCMV infection setting, the primed CD8 T cells first appear in the crypts and over subsequent days accumulate along the length of the villi, with cells near the tips having a more mature Trm gene expression profile.^20^

While chemokine receptors that couple to Gαi-containing heterotrimeric G-proteins have well established roles in guiding immune cell distribution, there have been increasing examples of Gα13-coupled receptors influencing cell positioning and homeostasis within tissues.^21–24^ A key downstream effector of Gα13 is Arhgef1, a guanidine nucleotide exchange factor (GEF) for Rho ^25,26^. Although Gα13 is closely related to Gα12, multiple studies have established non-redundant roles for these Gα-proteins. A diverse set of GPCRs can couple to Gα13 and in most immune cell studies, these receptors inhibit migration towards chemoattractants.^21–23,27^ A notable exception to this paradigm has been GPR132 (G2A), a receptor that can be stimulated in vitro by exposure to several lipid mediators including lysophosphatidylcholine (LPC), where a promigratory response occurs.^28–31^ So far, the impact on IEL of deficiency in one Gα13-coupled migration-inhibitory receptor, GPR55, has been reported and found to cause a mild increase in γδT cells.^21^ Whether other Gα13 coupled receptors influence intestinal T cell function is unclear.

Here we sought to broadly test the importance of Gα13-coupled receptors in intestinal immune homeostasis by deleting Gα13 in all hematopoietic cells. High dimensional flow cytometry revealed a striking loss of intestinal T cells that encompassed γδ T cells, CD8αα and CD8αβ αβT cells and that was T cell intrinsic. Conditional deletion of Gα13 led to a progressive loss of IEL that was preceded by a reduction in motility and increased apoptosis. Using a *Listeria monocytogenes (Lm)*-OVA intestinal infection model, Gα13-pathway deficient OVA-specific OT1 T cells reached the intestine in near normal numbers but failed to accumulate in the Trm compartment and showed reduced Bcl2 and deficient induction of TGFβ-pathway genes. Over-expression of Bcl2 or activated TGFβ was sufficient to partially restore Gα13-pathway deficient IEL. GPR132 was identified as a Gα13-coupled receptor contributing to CD8αβ IEL homeostasis. Finally, the lack of IEL caused by Gα13-deficiency led to more severe dextran sodium sulfate (DSS)-indued colitis and less ability to control colorectally implanted tumor cells.

## RESULTS

### Gα13 is required for intestinal IEL homeostasis

To broadly examine the role of Gα13 in intestinal immune cell homeostasis, Gna13f/f mice that had been crossed with the pan-hematopoietic VavCre line (Gα13^ΔVav^) were studied. Flow cytometric analysis with a 35 color antibody panel revealed a striking reduction in the CD8 T cell subsets with minimal impact on CD4 T cells (Fig. 1A). Analysis of further mice using a focused panel for CD8 and CD4 αβ and γδ T cells revealed a ∼100 fold deficiency in γδT and CD8αα αβT IEL, a ∼10-fold deficiency in CD8αβ αβT IEL, and a ∼5-fold deficiency in CD4 CD8α double positive αβT IEL, while CD4 αβT IEL were not significantly affected (Fig. 1B, C, S1A). Similar findings were made for the colon (Fig. 1D). Small intestinal lamina propria (LP) CD8 T cell subsets were also greatly reduced (Suppl. Fig. S1B). The marked deficiency in γδT IEL and CD8αβ IEL was confirmed in immunofluorescence microscopy of small intestinal tissue sections (Fig. 1E and Fig. S1C).

**Figure 1:**
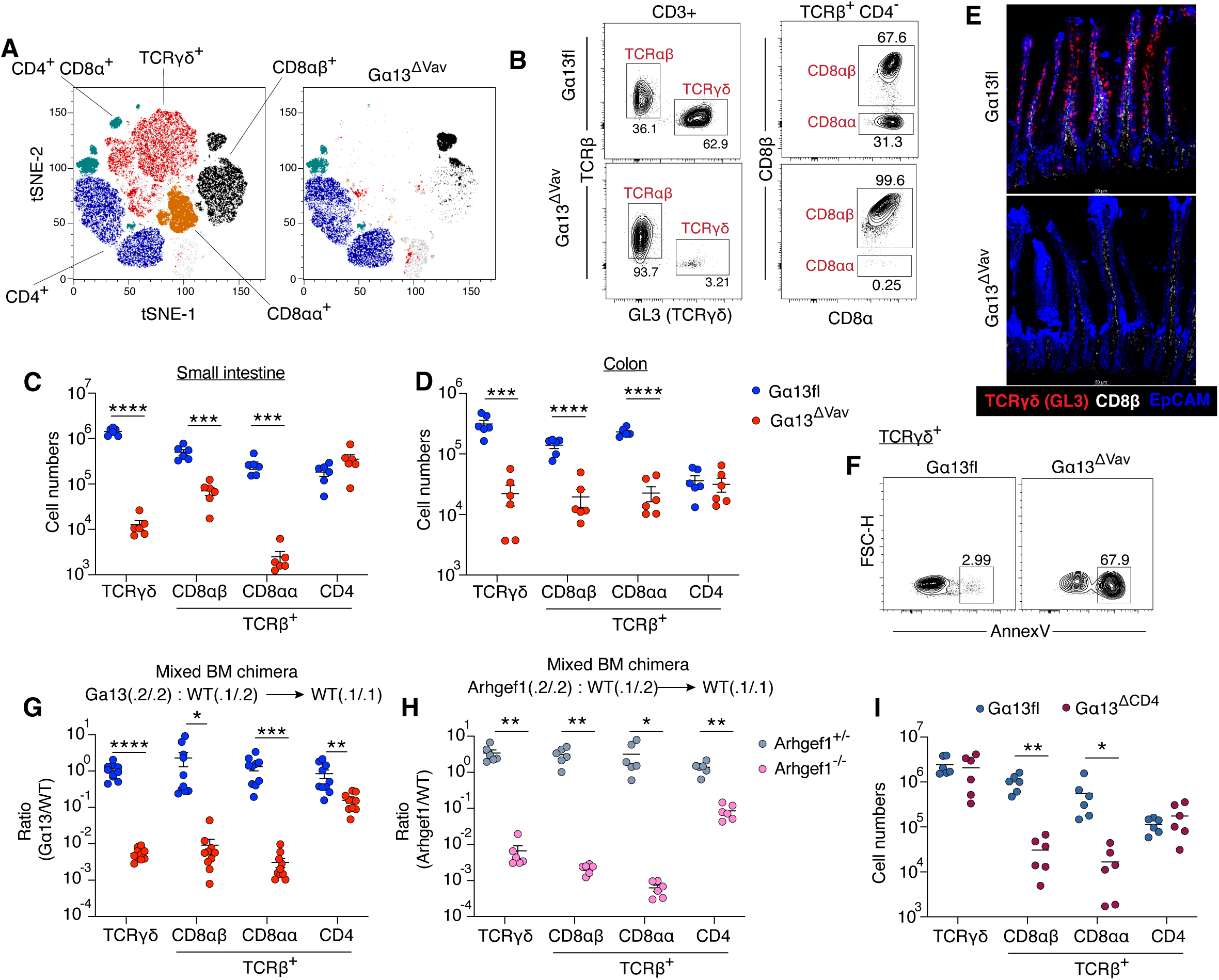
Gα13 is required for intestinal IEL T cell homeostasis. A. t-SNE analysis of 35 color flow cytometry panel (Table S1) of the small intestine from Gα13fl (left) and Gα13^ΔVav^ (right) mice. B. Representative flow cytometry plot of the frequency of small intestinal intraepithelial TCRαβ^+^ and TCRγδ^+^ cells among CD3^+^ cells (left) and TCRαβ^+^ CD8αβ^+^ or TCRαβ^+^ CD8αα^+^ among CD8α^+^ CD4^-^ cells (right) from Gα13fl and Gα13^ΔVav^ mice. C, D. Total small intestine (C) or colon (D) cell numbers of IEL subsets in Gα13fl and Gα13^ΔVav^ mice. (N= 6 mice/group) E. Representative IF image from the small intestine of Gα13fl and Gα13^ΔVav^ mice stained for TCRγδ (red), CD8β (white), and EpCAM (blue). F. Representative flow cytometry plot of the Annexin V^+^ cell frequency among small intestine TCRγδ^+^ cells from Gα13fl and Gα13^ΔVav^ mice G. Mixed BM chimera analysis for indicated small intestine IEL types showing ratios of CD45.2^+^ Gα13fl (blue) or Gα13^ΔVav^ (red) and CD45.1/2^+^ WT cells in WT recipients. (N= 10 mice/group). H. Same as in G except with CD45.2^+^ Arhgef1^+/-^ (grey) and Arhgef1^-/-^ (pink) with CD45.1/2^+^ WT. (N= 6 mice/group). I. Total numbers of small intestinal IEL subsets in Gα13fl and Gα13^ΔCD4^ mice. (N= 6 mice/group). All data are pooled from two independent experiments and represented as mean ± SEM. \*\*\*\**P*<0.0001, *** *P*<0.001, ** *P*<0.01, * *P*<0.05, t-test.

Intravascular labeling established that the few remaining IEL in Gα13^ΔVav^ mice were extravascular (Fig. S1D, E). γδT and CD8αβ T cell numbers in other non-lymphoid tissues (skin, liver, lung) and in spleen and lymph nodes (LNs) were not affected by Gα13-deficiency (Suppl. Fig. S1F, G). The number of thymic CD8αα IEL precursor cells^16^ was slightly increased making it unlikely that the loss in CD8αα IEL reflected reduced development (Suppl. Fig. S1H). Staining for surface phosphatidylserine, a marker of apoptosis, with Annexin V showed that most of the remaining γδT IEL in Gα13^ΔVav^ mice were Annexin V^+^ (Fig. 1F and Suppl. Fig. S1I).

Given the impact of Gα13-deficiency appeared strongest on CD8αα^+^ IEL, we also stained cells with the noncanonical class I molecule, TL, that binds CD8αα.^32^ This analysis revealed that the CD8αα TCRαβ cells remaining in Gα13^ΔVav^ mice maintained a similar ability to bind TL as WT cells in accord with their similar CD8αα expression (Fig. S1J). Interestingly, TL binding by CD8αβ IEL was reduced (Fig. S1J) as was the frequency of CD8αα+ γδ T cells (Fig. S1K).

To test for a T cell intrinsic role of Gα13 in IEL homeostasis we reconstituted irradiated mice with a mixture of WT (CD45.1/2) and Gα13^ΔVav^ or WT control (CD45.2) bone marrow (BM). Analysis of the mixed BM chimeras after 8 weeks of reconstitution showed a marked γδT, CD8αα αβT and CD8αβ αβT IEL deficiency, establishing the essential role of this Gα protein in CD8 IEL (Fig. 1G). Under these competitive conditions there was a reduction in Gα13-deficient CD4 IEL (Fig. 1G). Analysis of mice lacking the Gα13 effector, Arhgef1, showed a similar γδT and CD8 αβT IEL deficiency (Suppl. Fig. S1L). Using the mixed BM chimera approach, a similarly severe IEL defect was observed for Arhgef1-deficient as for Gα13-deficient chimeras (Fig. 1H) providing evidence that Gα13 acts in IEL homeostasis through engagement of Arhgef1.

To further test the αβT cell intrinsic role of Gα13 we generated Gna13f/f CD4Cre mice. CD4Cre is expressed in CD4 CD8 double positive (DP) thymocytes and therefore targets CD8 and CD4 T cells but not γδT cells.^33^ These mice showed a marked deficiency in CD8αα and CD8αβ αβT IEL, while γδT IEL were unaffected (Fig. 1I). Many CD8αβ T cells in WT mice expressed IFNγ as expected^34^ whereas few of the CD8αβ IEL remaining in Gα13^ΔCD4^ mice expressed IFNv (Suppl. Fig. S1M). Overall, these data establish a selective high dependence of intestinal CD8 IEL on Gα13- and Arhgef1-signaling.

### Conditional Gα13 deletion from IEL leads to defective motility and cell loss

To gain insight into the temporal requirement for Gα13 by IEL we crossed Gna13f/f mice with TCRδ-CreERT2 (Gα13^ΔTCRδ^) and E8iCreERT2 (Gα13^ΔCD8α^) mice to enable tamoxifen mediated deletion of the Gna13 gene in mature γδT cells and CD8 T cells, respectively. Based on prior evidence that the TCRδ-CreERT2 had low activity and deletion was enhanced in mice harboring two alleles^35^ we studied TCRδ-CreERT2 homozygous mice and we included an Ai6 Rosa26-LSL-zsGreen reporter to mark cells that had experienced Cre activity (Suppl. Fig. S2A). We used five consecutive doses of tamoxifen as in previous work with this mouse line.^35^ Three days following tamoxifen treatment, a time point when there had likely been only partial loss of Gα13, there had been little change in γδT IEL numbers (Fig. 2A). By 2 weeks there was a substantial loss of γδT IEL (Fig. 2A). The E8iCreERT2 is efficacious^36^, and we found strong induction of reporter (Ai14-tdTom) expression following three days of tamoxifen treatment (Suppl. Fig. S2B). Gα13-deleted CD8αα and CD8αβ αβT IEL decayed from the intestine over a 1 week period (Fig. 2B). E8iCre is also active in CD8^+^ γδT cells, and γδT IEL decayed over 1 week after tamoxifen treatment (Suppl. Fig. S2C).

**Figure 2:**
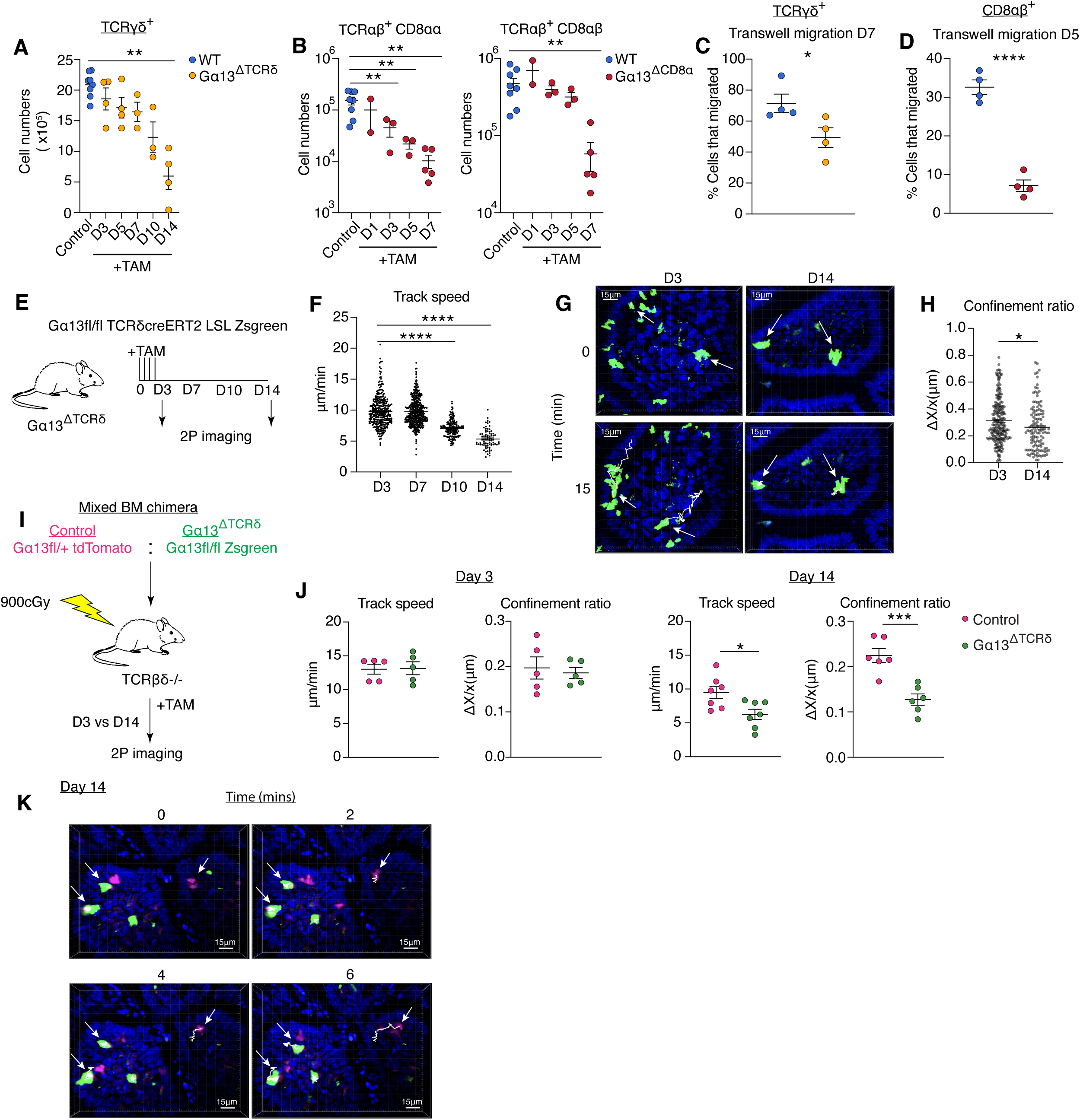
Conditional Gα13 deletion from IEL leads to cell loss and defective motility. A. Number of TCRγδ IEL in the small intestine of WT mice (blue, Gα13^ΔTCRδ^ – TAM) or inducible Gα13^ΔTCRδ^ mice (yellow, Gα13^ΔTCRδ^ + TAM) 3, 5, 7, 10 or 14 days after TAM treatment. (N=3-8 mice/group) B. Number of TCRαβ^+^ CD8αα^+^ (left) or TCRαβ^+^ CD8αβ^+^ (right) IEL in the small intestine from WT mice (blue, Gα13^ΔCD8α^ – TAM) or inducible Gα13^ΔCD8α^ mice (maroon, Gα13^ΔCD8α^ + TAM) 1, 3, 5, 7 or 14 days after TAM treatment. In A and B, data are from three independent experiments. (N=2-8 mice/group) C,D. Frequency of TCRγδ^+^ (C) or TCRαβ^+^ CD8αβ^+^ (D) IELs that migrated through a transwell without added ligands 7 or 5 days, respectively, after TAM treatment. Groups and coloring as in (A,B). (N=4 mice/group) E. Experimental scheme of two-photon intravital imaging of the small intestine of Gα13^ΔTCRδ^ ZsGreen mice 3, 7, 10, and 14 days after TAM treatment. F. Track speed (µm/min) of ZsGreen^+^ TCRγδ cells in mice treated as in E. Data are pooled for all cells imaged in a field of view from N=3 mice imaged at each time point. G. Time series at day 3 and day 14 showing representative tracks at 0 and 15 minutes of imaging. White arrows highlight cells with marked tracks (white lines). See also Movies S1-S4. H. Confinement ratio of TCRγδ cells (displacement/distance) determined from the tracks of cells at D3 or D14 after TAM. Data are pooled for all cells imaged in a field of view from N=3 mice imaged at D3 vs D14. I. Experimental scheme of mixed BM chimera two-photon intravital imaging of TCRγδ cells 3 or 14 days after TAM treatment. TCRβδ^-/-^ recipient mice were irradiated and reconstituted with control BM (pink, Gα13fl/+ TCRdCreERT2^+^ tdTomato) mixed 1:1 with Gα13^ΔTCRδ^ BM (green, Gα13fl/fl TCRdCreERT2^+^ ZsGreen). J. Track speed (µm/min) and confinement ratio plotted as the mean for all cells of each type tracked in each mouse at day 3 or 14 after TAM from mixed BM chimeras of the type in I. (N= 5 mice/group). K. Representative tracks over time (0, 2, 4, 6 min) of control (pink) and Ga13^ΔTCRδ^ (green) TCRγδ cells 14 days after TAM. White arrows highlight cells with marked tracks (white lines). See Movies S5 and S6. All graphed data are pooled from two independent experiments and represented as mean ±χσβαρλινε SEM. \*\*\*\**P*<0.0001, *** *P*<0.001, ** *P*<0.01, * *P*<0.05, t-test.

We next asked how Gα13-deficiency impacted IEL motility. In transwell assays, T cells from lymphoid organs show very limited migration (<5%) in the absence of chemokine.^37,38^ In agreement with earlier work^21^ we found that γδT IEL and CD8αβ T IEL had high basal migration in the absence of chemokine (30-70%). This constitutive motility was strongly dependent on Gα13 (Fig. 2C, D). To test the impact of Gα13-deficiency on IEL migration in vivo, we used intravital 2-photon microscopy.^2,3^ Intravital imaging of Gα13^ΔTCRδ^ Ai6-ZsGreen reporter mice at day 3 after tamoxifen treatment (Fig. 2E), when the reporter had been activated but Gα13 protein likely had minimally decayed, revealed track speeds (Fig. 2F) and migration dynamics (Fig. 2G, Movie S1) similar to previous reports.^2,3,39^ By day 7, when γδT cell numbers had not significantly declined, the track speed and migration behavior was not significantly changed (Fig. 2G and Movie S1). However, by day 10 the track speed had decreased and for the reporter^+^ cells remaining at day 14 there was a striking reduction in migration with some cells moving and then stopping in association with the epithelium for the remaining duration of the movie (Fig. 2G and Movie S1). The confinement ratio, measured as displacement divided by track length, was significantly reduced at day 14 (Fig. 2H). In a second approach, to allow imaging of WT and Gα13-deficient IEL in the same tissue, irradiated TCRβδ^-/-^ mice were reconstituted with Ai14-tdTomato control and Gα13^ΔTCRδ^ Ai6-ZsGreen BM. After an ∼8 week reconstitution period, mice were treated with tamoxifen and the intestine imaged at day 3 and day 14 (Fig. 2I). At day 3 after tamoxifen both control γδT cells and Gα13-deficient γδT cells were highly motile (Movies S2). However, 14 days after tamoxifen, the control γδT cells remained motile while most Gα13-deficient γδT cells in the same tissue volume showed greater confinement, failed to move or arrested their migration during the imaging period (Fig. 2J-L, and Movie S2). Taken together, these findings provide evidence that Gα13 expression contributes to the intrinsic motility of IEL and is required for their epithelial patrolling behavior.

### Gene expression profile of Gα13-deficient intestinal T cells

To gain insight into the impact of Gα13-deficiency on IEL gene expression we performed 10x scRNAseq on a pool of sorted TCRγδ^+^ and TCRαβ^+^ CD8α^+^ and CD4^+^ IEL from control and Gα13^ΔVav^ mice. This analysis revealed a striking shift in the gene expression profile of the γδT and CD8αβ αβT IEL remaining in Gα13^ΔVav^ mice (Fig. 3A and Suppl. Fig. S3 A,B). The CD8αα αβT IEL were not well represented due to their several fold lower frequency amongst CD8α^+^ TCRαβ^+^ IEL than CD8αβ^+^ cells (Fig. 1C). An increased fraction of both γδT and αβT IEL were proliferating, perhaps a consequence of the marked loss of these cell types leaving an open niche that over-stimulates the rare survivors. The CD4 T cells also showed altered gene expression (Fig. 3A) possibly due to the profound loss of CD8^+^ IEL but perhaps also reflecting intrinsic changes as suggested by their reduced competitiveness in mixed BM chimeras (Fig. 1G). Dot plot analysis showed reduced integrin *Itgae* and *Itga1* expression in γδT and CD8αβ T IEL and reduced expression of IEL-associated transcription factors *Nr4a2*, *Nr4a3*, *Junb*, and *Fosb* and of the Trm transcription factor *Runx3*. Notably, γδT cells had markedly reduced *Bcl2* expression and there was increased expression of pro-cell death genes *Zbp1*, *Casp3*, *Casp4*, *Bax* and *Fas* (Fig. 3B). The Gα13-deficient cells also had higher expression of certain genes associated with naïve cells, including *Klf2*, *S1pr1* and *Ly6a*, perhaps indicative of a less differentiated state and recent recruitment to the IEL compartment (Fig. 3B). Another notable feature of the Gα13-deficient γδT cells was a reduction in an IL15-induced signature ^40^ (Fig. 3C). A concern with this analysis was that the broad loss of Gα13 in hematopoietic cells may indirectly influence gene expression in the remaining IEL. We therefore performed RNAseq on CD8αβ IEL sorted from Gα13^ΔCD4^ mice. Among top genes that were more highly expressed in Gα13-deficient CD8αβ IEL were *Tcf7*, *Klf2*, *S1pr1*, *Ccr7* and *Sell*, genes that are highly expressed in naïve T cells, again suggesting a less differentiated state (Fig. 3D). Genes that were less expressed in KO cells included effector program associated molecules such as *Prdm1*, *Itga1*, *Gzma*, the cytokine *Ifng*, IEL-associated transcription factors *Egr2*, *Nr4a2*, *Nr4a3*, *Junb*, and *Fos*, and survival promoting factors *Il7r* and *Bcl2* (Fig. 3D, E). Pathway analysis highlighted impacts of Gα13-deficiency on chemotaxis/migration, TGFβ, STAT5 and NFkB signaling, mTORC signaling, and programmed cell death (Fig. 3E).

**Figure 3:**
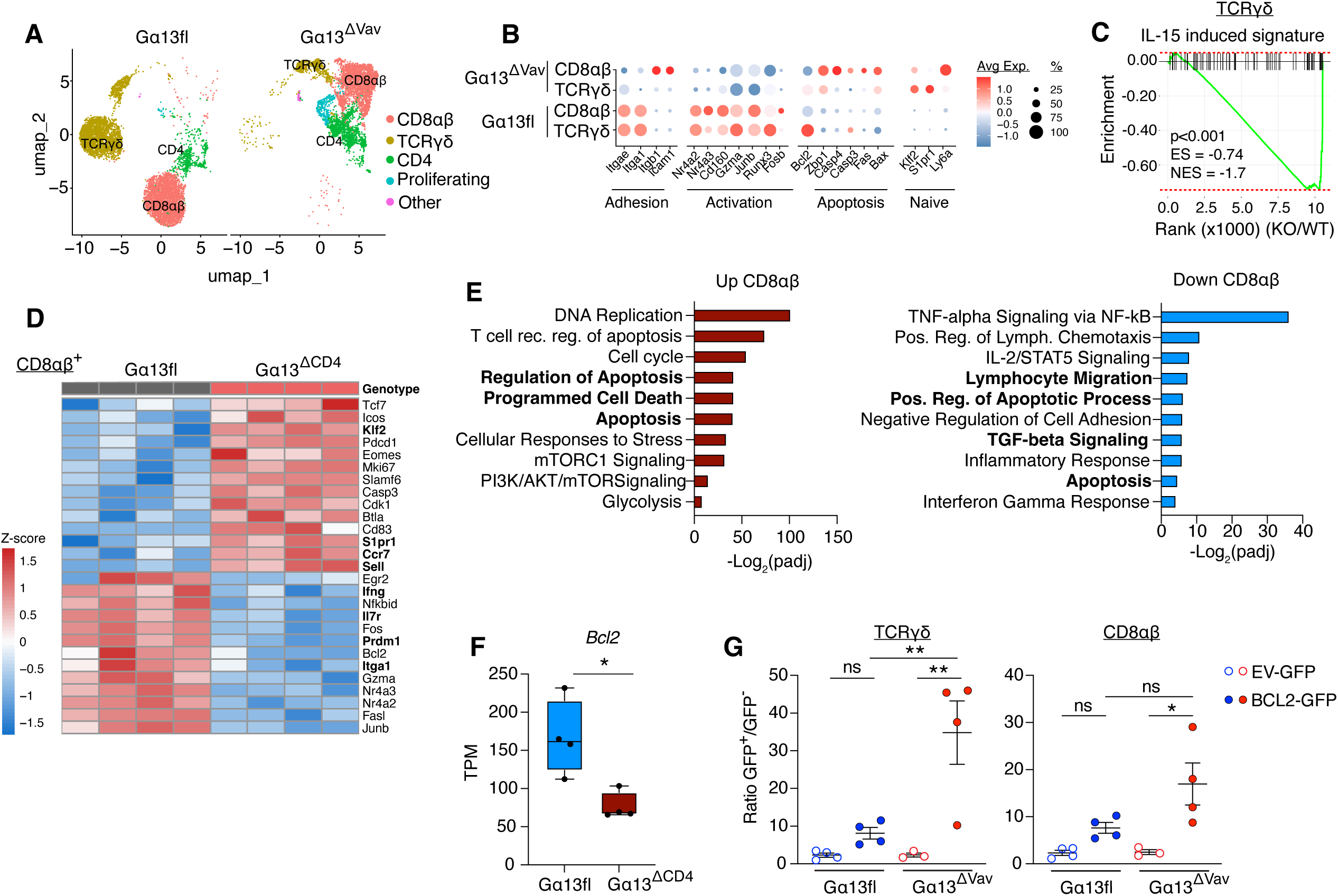
Gene expression profile of Gα13-deficient intestinal T cells. A. uMAP plot of 10X scRNA-seq data of matched numbers of IELs sorted from Gα13fl (left) and Gα13ΔVav (right) mice. B. Dot plot of select DE genes from CD8αβ and TCRγδ T cell clusters of Gα13fl and Gα13^ΔVav^ mice. C. GSEA of IL15-induced signature ^40^ in TCRγδ cells from Gα13^ΔVav^ mice compared to Gα13fl mice. D. Heatmap of select DE genes from SMARTseq of CD8αβ IELs from Gα13fl and Gα13^ΔCD4^ mice. E. Pathway analysis for (left) upregulated and (right) downregulated genes, showing select differently expressed biological pathways from MsigDB hallmark, Gene Ontology, WikiPathways, and KEGG in CD8αβ IELs from Gα13fl and Gα13^ΔCD4^ mice. Padj<0.05. F. Expression of *Bcl2* (TPM, transcripts/million) in CD8αβ IELs from Gα13fl and Gα13^ΔCD4^ mice. (N=4 mice/group) G. Retroviral BM chimeras with overexpression of EV-GFP (open circles) or Bcl2-GFP (filled circles) in Gα13fl (blue) and Gα13^ΔVav^ (red) cells. Data are expressed as a ratio of GFP^+^/GFP^-^ of each group of TCRγδ (left) or CD8αβ (right) IELs. N = 3-5 mice/group and are representative of two independent experiments. Data in F, G are pooled from two independent experiments and shown as mean ± SEM. * *P*<0.05, t-test (F), or ** *P*<0.01, * *P*<0.05, ANOVA with Tukey multiple comparisons (G).

The reduced *Bcl2* expression in γδT and CD8αβ αβT IEL (Fig. 3B, D, F) together with the evidence for increased apoptosis (Fig. 1F) led us to test the sufficiency of Bcl2 over-expression to rescue Gα13-deficient IEL. BM from control (Gα13fl) or Gα13^ΔVav^ mice was transduced with a retroviral construct encoding Bcl2 and a GFP reporter, or an empty vector encoding only GFP (EV-GFP). Analysis of the intestine of the chimeric mice 8 weeks after reconstitution showed Bcl2 was sufficient to greatly increase γδT and CD8αα and CD8αβT IEL in Gα13^ΔVav^ mice (Fig. 3G and Fig. S3C). These findings provide evidence that Gα13 function is required for intact signaling by several pathways and has effects that promote IEL survival.

### Gα13 signaling controls the maturation and homeostasis of intestinal CD8αβ Trm

To test the role of G13-pathway signaling during intestinal CD8αβ IEL induction, we employed the oral *Lm*-OVA infection and OTI TCR transgenic adoptive cell transfer model.^18^ We used Arhgef1-deficient mice for these studies as the full genetic knockout avoided concerns about incomplete deletion of a floxed allele. Control or Arhgef1-deficient OTI cells were transferred into mice that were gastrically infected with *Lm*-OVA (Fig. 4A). Analysis 8 days later revealed similar intestinal recruitment of control and Arhgef1-deficient cells (Fig. 4B). However, the number of Arhgef1-deficient cells began decaying relative to control by day 11 and they were 100-fold depleted by day 28 (Fig. 4B). Annexin V staining showed higher apoptosis amongst Arhgef1-deficient OTI cells was already evident at day 8 (Fig. 4C).

**Figure 4.**
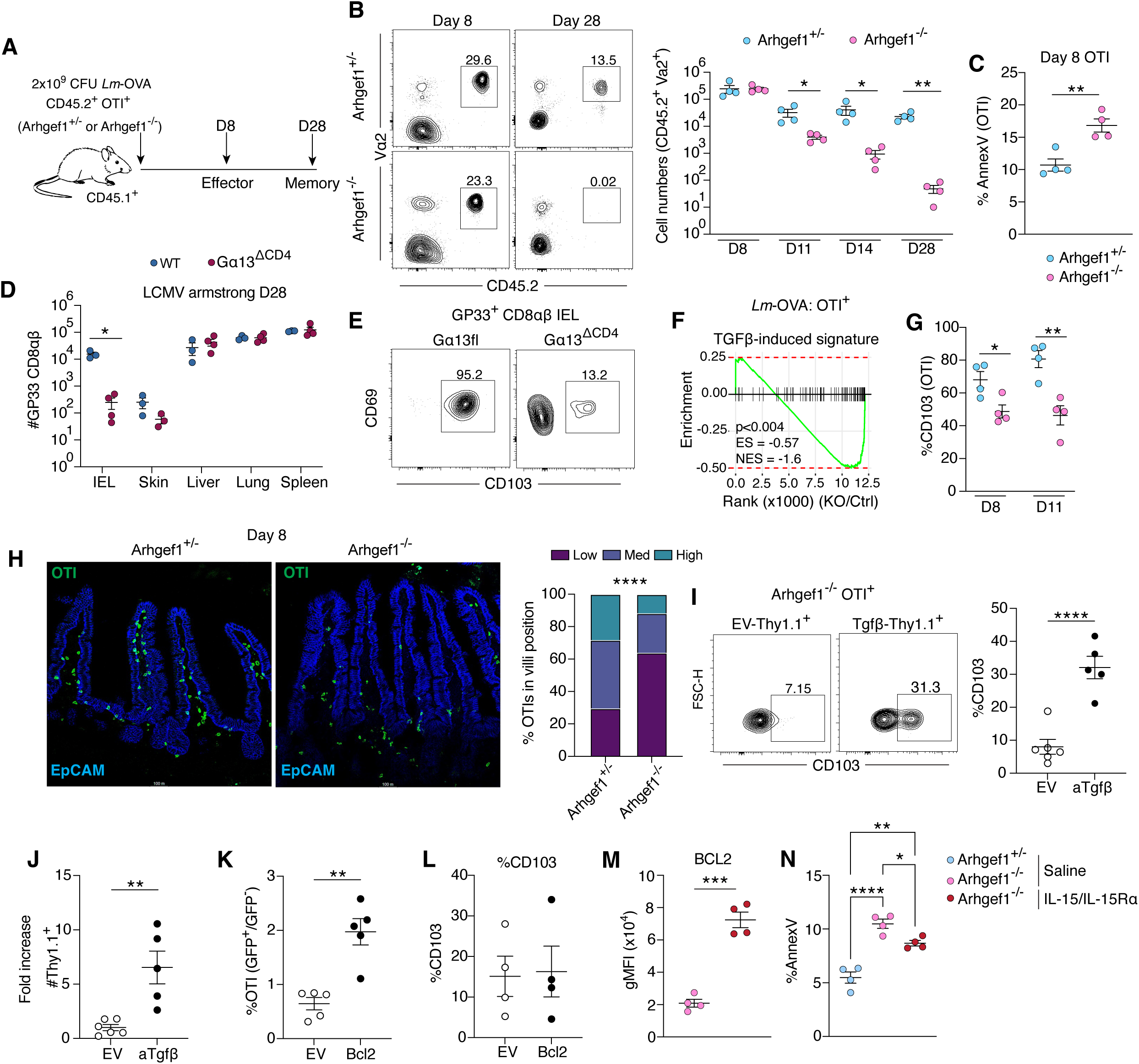
Gα13 signaling controls the maturation and homeostasis of intestinal CD8αβ Trm. A. Experimental scheme for analyzing effector vs memory intestinal Arhgef1^+/-^ or Arhgef1^-/-^ OTI T cell response following adoptive transfer and oral *Lm*-OVA infection. B. Left: representative FACS plots of Arhgef1^+/-^ and Arhgef1^-/-^ OTI (Vα2^+^ CD45.2^+^) T cells among CD8αβ IEL 8 or 28 days after infection. Right, absolute cell numbers of Arhgef1^+/-^ and Arhgef1^-/-^ OTI T cells in small intestine epithelium 8, 11, 14, and 28 days after infection. (N=4 mice/group) C. Frequency of Annexin V^+^ Arhgef1^+/-^ or Arhgef1^-/-^ OTI (Vα2^+^ CD45.2^+^) T cells by FACS 8 days after infection. (N=4 mice/group) D. Number of GP33^+^ CD8αβ^+^ T cells 28 days after LCMV Armstrong infection of WT (or Gα13^ΔCD4^ mice (maroon) in small intestine (IEL), skin, liver, lung, and spleen. (N=3-4 mice/group) E. Representative FACS plot of CD69^+^ and CD103^+^ GP33^+^ CD8αβ Trm cells in the IEL from WT or Gα13^ΔCD4^ mice. F. GSEA of TGFβ induced signature (Reina-Campos et al. 2025) in CD8αβ T cells comparing Arhgef1^+/-^ and Arhgef1^-/-^ OTI T cells at day 8 of the *Lm-*OVA infection time course. G. Frequency of CD103^+^ Arhgef1^+/-^ and Arhgef1^-/-^ OTI T cells 8 or 11 days after infection. (N=4 mice/group) H. Left panels show representative IF images of small intestinal sections from recipients of Arhgef1^+/-^ or Arhgef1^-/-^ OTI T cells (CD45.2, green) at day 8 after infection, EpCAM (blue). Right graph shows summary frequency of OTI T cells found in the low, med, and high position of the villi. (N=4 mice/group). I. Retroviral overexpression of aTGFβ-Thy1.1 and control EV-Thy1.1 in Arhgef1^-/-^ OTI T cells. Left panels are representative flow plots showing frequency of CD103+ in Thy1.1^+^ OTI T cells for EV and aTGFβ groups at day 11 post infection. Right graph shows summary CD103 data. (N=5-6 mice/group). J. Fold increase in the absolute number of reporter+ cells from (I). K. Retroviral overexpression of Bcl2-GFP or control EV-GFP in Arhgef1^-/-^ OTI T cells. The cells were mixed 1:1 with non-transduced Arhgef1^-/-^ OTI T cells, transferred and analyzed at day 11 post infection. (N=5 mice/group). L. Frequency CD103^+^ among GFP^+^ EV and Bcl2 OT1 cells from (K). (N=5 mice/group). M. Geometric mean fluorescence intensity of BCL2 in Arhgef1^-/-^ OTI cells analyzed at day 8 of infection and day 2 of treatment with saline or IL15/IL15Ra. (N=4 mice/group) N. Frequency of AnnexinV^+^ Arhgef1^+/-^ or Arhgef1^-/-^ OTΙ cells analyzed at day 8 of infection and day 2 of treatment with saline or IL15/IL15Ra. (N=4 mice/group). All data are pooled from two independent experiments and represented as mean ± SEM. \*\*\*\**P*<0.0001, *** *P*<0.001, ** *P*<0.01, * *P*<0.05, t-test. H, \*\*\*\**P*<0.0001, Fisher’s exact test, N, *** *P*<0.001, ** *P*<0.01, * *P*<0.05 ANOVA with Tukey multiple comparison test.

We took advantage of the OTI transfer system to confirm that Gα13 and Arhgef1 were acting in the same pathway. Control and Arhgef1-deficient OTI cells were electroporated with Cas9-sgRNA ribonucleoprotein (RNP) complexes to inactivate Gα13 prior to adoptive transfer. The Gna13 targeting RNPs had an ∼80% mutation rate and they were sufficient to cause a significant loss of OTI cells compared to control targeted cells (Fig. S4A). However, mutation of Gna13 in Arhgef1-deficient cells did not have any additive effect, in accord with these genes acting in the same pathway (Fig. S4A).

To determine the intestinal selectivity of the G13-pathway dependence, we systemically infected WT and Gα13^ΔCD4^ mice with LCMV Armstrong, a pathogen that infects most organs of the mouse and broadly induces CD8 Trm formation.^11^ We assessed LCMV-specific CD8 Trm in tissues using Gp33 tetramers at day 28. There was a marked reduction in tetramer^+^ IEL in the intestine but little reduction in skin, lung, spleen or LNs (Fig. 4D). The few remaining GP33^+^ CD8αβ IELs in Gα13^ΔCD4^ mice at day 28 revealed a striking defect in tissue resident memory markers CD69 and CD103 (Fig. 4E). Although the appearance of similar numbers of OTI CD8 T cells in the intestine at day 8 indicated intact homing, to further test this we activated WT and Gα13 KO CD8 T cells in vitro in the presence of RA and TGFβ to promote induction of CCR9 and α4β7 ^41^. The Gα13-deficient cells proliferated equivalently to WT cells, showed equivalent upregulation of CCR9 and α4β7, and intact transwell migration to CCL25 (Suppl. Fig. S4B, C). Adoptive transfer of these polyclonal gut tropic CD8 T cells to WT mice showed similar homing to the small intestine (Suppl. Fig. S4D). However, after 5 days, there were less Gα13-deficient compared to WT cells and by day 10 the loss of Gα13-deficient cells was marked (Suppl. Fig. S4D).

Based on RNAseq analysis of day 8 transferred OTI cells, there was reduced TGFβ pathway activity in the absence of Arhgef1 (Fig. 4F). This included reduced expression of *Itgae* that encodes integrin αE, also called CD103, a known TGFβ-induced gene. Reduced CD103 expression was confirmed by flow cytometry and occurred at day 8 in the tissue, before notable cell loss was observed (Fig. 4G). Control OTI cells upregulated CD103 from day 8 to day 11, however this process was impaired in the Arhgef1-deficient cells in accord with a defect in TGFβ signaling and thus Trm differentiation in the tissue (Fig. 4G). Gα13-deficient CD8 T cells demonstrated an intact ability to upregulate CD103 in response to TGFβ in vitro making it unlikely that Gα13 was directly involved in TGFβR signaling (Fig. S4E). Immunofluorescence microscopy of intestinal tissue at day 8 after transfer showed control OTI cells were distributed along the length of intestinal villi whereas Arhgef1-deficient OTI were often more enriched near the villus base (crypt) (Fig. 4H and Suppl. Fig. S4F). This distribution is similar to findings for TGFβR^-/-^ P14 CD8αβ T cells at the same time point.^20^

To test whether the reduced TGFβ-induced genes contributed to the CD103 deficiency and impaired IEL retention, we transduced Arhgef1-deficient OTI cells with empty vector (EV) or an activated TGFβ (aTGFβ) construct^42^ prior to adoptive transfer to *Lm*-OVA infected recipients.

The activated TGFβ has a mutation that limits association of the TGFβ small latent complex with the latent TGF binding protein-1 and tethering to the extracellular matrix ^42^. Expression of this construct by T cells is expected to at least partially overcome the normally tight spatial restriction of active TGFβ ^43^. Activated TGFβ over-expression partially rescued CD103 expression (Fig. 4I) and prevented the decay of Arhgef1-deficient OTI CD8αβ IELs from the tissue at day 11 (Fig 4J). The incomplete rescue of CD103 expression may indicate some continued restriction in TGFβ access but might also reflect an effect of the cell culture and retroviral transduction procedure on cell function, a possibility suggested by the lower CD103 expression by EV transduced versus untransduced OTI cells at day 8 after transfer (Fig. 4G and I). In contrast to the rescuing effect on Arhgef1-deficient cells, the ectopic TGFβ did not lead to increased generation of WT IEL likely due to WT cells already being well exposed to active TGFβ (Fig. S4G).

Beyond the effects of TGFβ, survival signals, such as IL15 or IL7, are needed to upregulate Bcl2 and prevent apoptosis.^11^ We therefore tested the impact of Bcl2 over-expression and observed a rescue in IEL frequency but no increase in CD103 expression (Fig. 4K, and L and Suppl. Fig. S4H). Bcl2 over-expression also did not rescue the high basal motility of Arhgef1-deficient IEL or their lack of distribution along the length of the villi (Fig. S4I, J). Moreover, in the *Lm*-OVA OTI transfer model, treatment of the mice with IL15/IL15Rα agonistic complexes^44^ rescued Arhgef1-deficient CD8αβ IEL Bcl2 expression and reduced the fraction of cells that were apoptotic without rescuing cell distribution (Fig. 4M, N, and Fig. S4K-N).Collectively these data suggest that Gα13 signaling is required for proper cell migration and positioning in the intestine for CD8αβ T cells to access niche-restricted cytokines, such as TGFβ and IL15, that support Bcl2 and CD103 upregulation and formation of long-lived Trm IELs.

### Gα13-coupled GPCR CRISPR screen identifies role for GPR132 in IEL homeostasis

Based on an assessment of RNAseq data from CD8αβ IEL for GPCR expression (Suppl. Fig. S5A), together with evidence curated from the literature regarding potential Gα13-coupling, we identified a set of 12 candidate GPCRs for in vivo testing. BM from Cas9-transgenic mice was transduced with retroviral constructs encoding pairs of sgRNA against individual GPCRs, Gna13 or control vectors (with Thy1.1 and BFP reporters), and used to reconstitute irradiated WT mice. Small intestinal IEL analysis of the 8 week reconstituted chimeras confirmed the efficacy of Gna13 targeting in reducing TCRαβ IEL (Fig. 5A). Of the 12 GPCRs, GPR132 targeting led to a significant reduction in IEL (Fig. 5A). Mixed BM chimeras generated using BM from GPR132^−/−^ ^45^ and WT mice showed reduced representation of GPR132-deficient CD8αβ T cells amongst IEL and LP cells, but not spleen cells (Fig. 5B, C and Suppl. Fig. S5B). The representation of TCRγδ IEL, CD8αα TCRαβ IEL, CD4 IEL and CD4 LP T cells were not significantly affected (Fig. S5B). In transwell migration assays, WT CD8αβ IEL migrated to LPC and this response was lost in GPR132-deficient IEL (Fig. 5D). Gα13-deficient CD8αβ IEL failed to migrate to LPC (Fig. 5E). The LPC promigratory activity on CD8αβ IEL was also Arhgef1-dependent (Fig. 5F).

**Figure 5.**
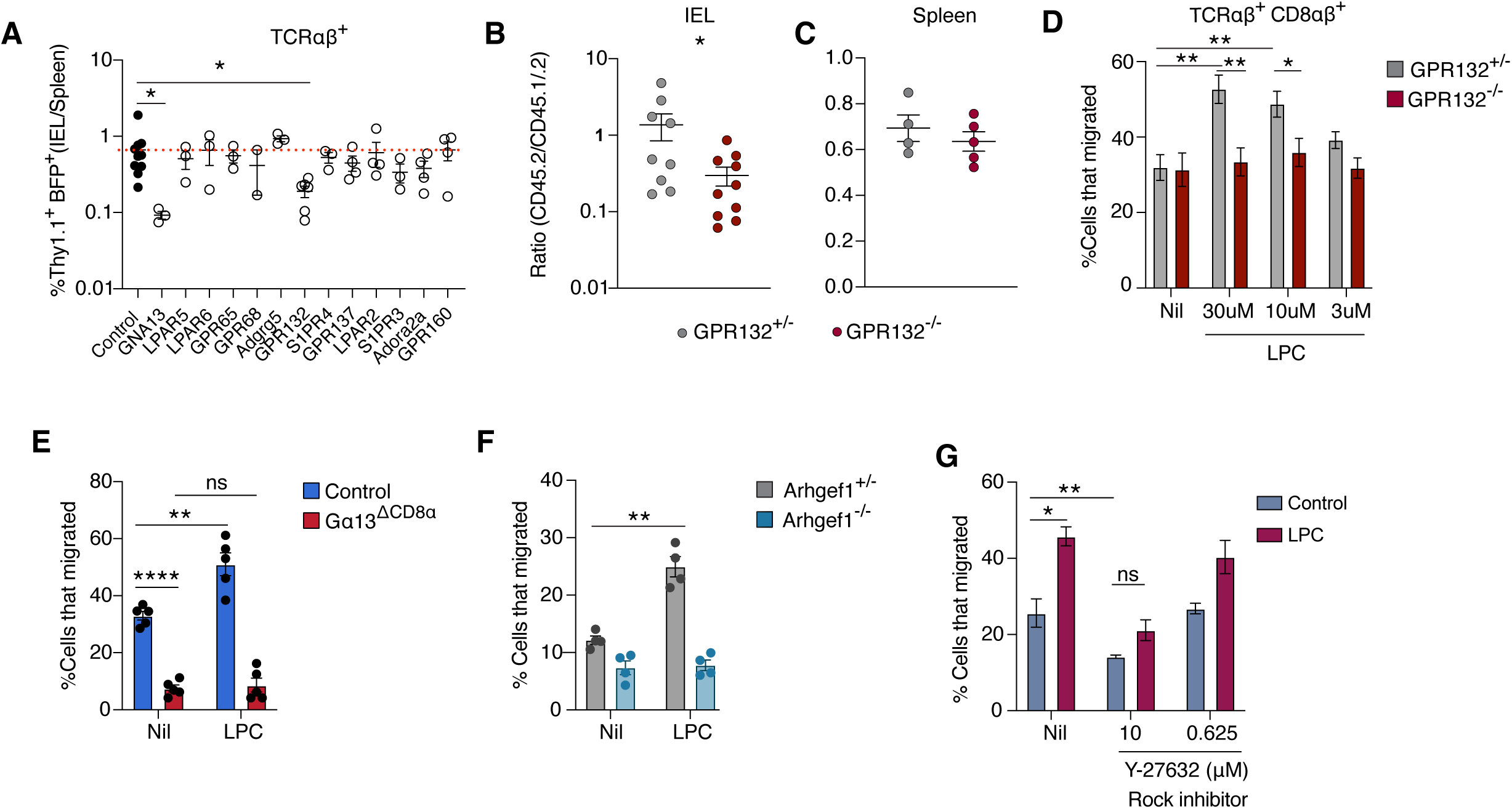
Gα13-coupled GPCR CRISPR screen identifies role for GPR132 in IEL homeostasis. A. Retroviral CRISPR/CAS9 BM screen of selected GPCRs. Data are expressed as a ratio of sgRNA1-BFP^+^ and sgRNA2-Thy1.1^+^ among TCRαβ^+^ cells in the IEL/spleen. Control represents non-targeting guides. (N=3-10 mice/group). B, C. Ratio of CD8αβ T cells in the IEL (B) or spleen (C) from mixed BM chimeras made with GPR132^+/-^ or GPR132^-/-^ (CD45.2) BM mixed 1:1 with WT (CD45.1/.2) BM. (N=4-9 mice/group). D. Frequency of GPR132^+/-^ or GPR132^-/-^ CD8αβ IELs that migrate in response to no ligand (Nil) or LPC added at 30, 10, or 3 µM to the bottom of a transwell. (N=4 mice/group). E. Frequency of CD8αβ IEL transwell migration to Nil, or LPC, of WT or inducible Gα13^ΔCD8α^ cells 5 days after TAM treatment. (N=5 mice/group). F. Frequency of CD8αβ IEL transwell migration to Nil or LPC of Arhgef1^+/-^ or Arhgef1^-/-^ OTI IEL at day 8 post *Lm-*OVA infection. (N=4 mice/group). G. Transwell CD8αβ IEL migration to LPC (maroon) or no ligand (grey blue) in the absence (Nil) or presence of ROCK inhibitor added at indicated concentrations. (N=4 mice/group). All data are pooled from two independent experiments and represented as mean ± SEM. \*\*\*\**P*<0.0001, *** *P*<0.001, ** *P*<0.01, * *P*<0.05, t-test.

The lower baseline motility in this experiment with transferred OT1 cells (Fig. 5F) compared to the experiments with endogenous IEL (Fig. 5G, E) may indicate that day 8 OT1 IEL have not yet acquired all properties of a mature IEL state. Arhgef1 facilitates the exchange of GDP to GTP for the small GTPase RhoA which activates the ROCK1/2 kinases.^25,26^ Treating CD8αβ IELs with the ROCK inhibitor, Y-27632, blocked migration in a dose dependent manner (Suppl Fig. S5C) and the ability of CD8αβ IELs to migrate to LPC (Fig. 5G). These findings provide evidence that GPR132 is a receptor acting upstream of Gα13, Arhgef1, and ROCK1/2 in CD8αβ IEL that contributes to their migration and competitive fitness. Nevertheless, GPR132 deficiency did not cause as severe a CD8αβ IEL loss as Gα13- or Arhgef1-deficiency. Three other Gα13 coupled receptors with known ligands, GPR55 and LPARs 5 and 6, were highly expressed by CD8αβ IELs (Suppl. Fig. S5A). Although CRISPR-targeting of these receptors did not lead to a loss of CD8αβ IELs alone, we observed their ligands, LPI and LPA, respectively, inhibited IEL migration in a Gα13 dependent manner (Suppl. Fig. S5D). It is therefore possible that combinations of these receptors signal through Gα13 to maintain the CD8αβ^+^ IEL populations.

### IEL loss due to Gα13 deficiency is associated with more severe colitis and reduced rectal tumor control

We next investigated the impact of Gα13-deficiency on susceptibility to inflammatory intestinal conditions. We first performed an intestinal permeability assay. WT or Gα13^ΔVav^ mice were gavaged with FITC-dextran and the serum concentration measured 4 hours later. We observed an increase in FITC-dextran permeability in the Gα13-deficient mice (Fig. 6A). To determine whether this altered barrier integrity contributes to bacterial translocation in inflammatory conditions, we infected mice with the gut restricted pathogen *Citrobacter rodentium*. We observed translocation of the pathogen to the MLN and spleen in Gα13^ΔVav^ mice but not control mice 10 days after infection (Fig. 6B). We next subjected Gα13-deficienct mice to dextran sodium sulfate (DSS) induced colitis. Following 5 days of exposure to 2% DSS in the drinking water, Gα13^ΔVav^ mice showed significantly greater weight loss (Fig. 6C), more intestinal pathology (Fig. 6D) and greater colon shortening compared to control mice (Fig. 6E). Similar weight loss findings were made with Gα13^ΔCD4^ mice (Fig. 6F) suggesting Gα13 signaling in T cells maintains the intestinal barrier during inflammation and protects the host from chemical-induced damage.

**Figure 6.**
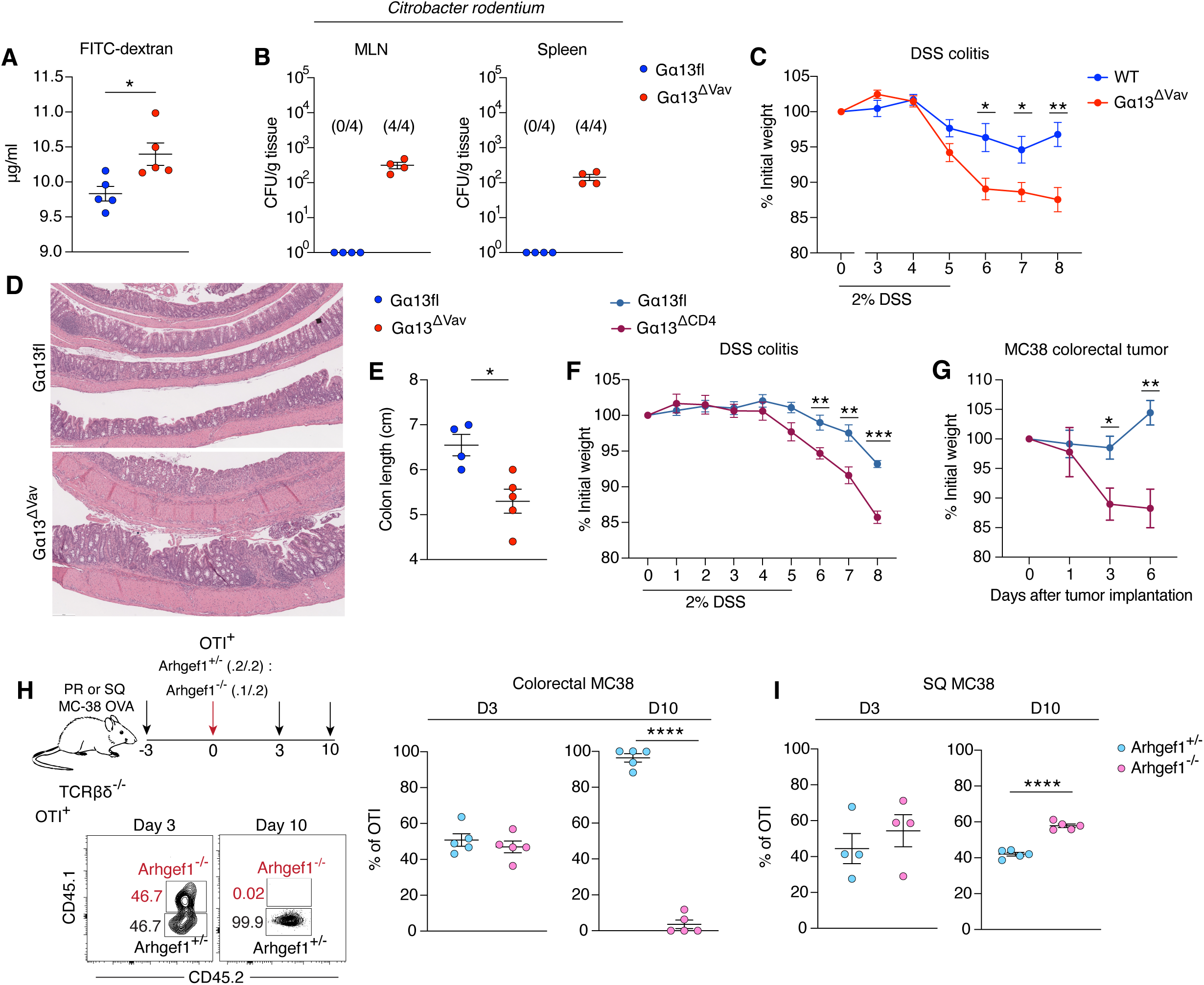
IEL loss due to T cell Gα13 deficiency is associated with more severe colitis and reduced rectal tumor control. A. Concentration (μg/ml) of FITC-dextran measured in the serum 4 hours after oral gavage of Gα13fl and Gα13^ΔVav^ mice. (N=5 mice/group). B. *Citrobacter rodentium* translocation to MLN or spleen 10 days after infection, shown as CFU/g tissue. (N=4 mice/group). C. Weight loss over time of Gα13fl and Gα13^ΔVav^ mice challenged with 2% DSS in the drinking water for 5 days. (N= 8-11 mice/group). D. Representative HCE from Swiss rolls of the colon of Gα13fl and Gα13^ΔVav^ mice 8 days after DSS. E. Colon length (cm) 8 days after DSS colitis in the indicated mice. (N=4-5 mice/group). F. Weight loss over time of Gα13fl and Gα13^ΔCD4^ mice challenged with 2% DSS in the drinking water for 5 days. (N= 8 mice/group). G. Weight loss over time of Gα13fl and Gα13^ΔCD4^ mice challenged with intrarectal MC38 tumors. (N=5-6 mice/group). H. Experimental scheme (top) of a mixed 1:1 Arhgef1^+/-^ (CD45.2) and Arhgef1^-/-^ (CD45.1/.2) OTI T cell transfer to TCRβδ^-/-^ mice that were intrarectally implanted with MC38-OVA tumors. Representative FACS plot (bottom) and summary data (right) of OTI T cells (vα2^+^ vβ5^+^) from the tumor 3 or 10 days after transfer, with Arhgef1 genotype identification based on CD45.1/.2 staining. (N=5 mice/group). I. Frequency of Arhgef1^+/-^ (CD45.2) and Arhgef1^-/-^ (CD45.1/.2) OTI cells transferred as in (H) from subcutaneously implanted MC38-OVA tumors analyzed 3 or 10 days after transfer. (N=4-5 mice/group). All data are pooled from two independent experiments and represented as mean ± SEM. \*\*\*\**P*<0.0001, *** *P*<0.001, ** *P*<0.01, * *P*<0.05, t-test.

To test the impact of Gα13-pathway deficiency in an intestinal tumor setting, we used the colorectally implanted MC38 tumor cell model.^46–48^ Gα13^ΔCD4^ mice suffered greater weight loss than littermate control mice following tumor implantation (Fig. 6G). Moreover, the Gα13^ΔCD4^ mice harbored larger tumors (Fig. S6A). To investigate tumor specific CD8 T cell responses, we rectally implanted MC38 cells expressing OVA in T cell-deficient mice and transferred an equal mixture of CD45.2 Arhgef1^+/-^ OTI and CD45.1/.2 Arhgef1^-/-^ OTI cells. Transferred control and Arhgef1-deficient OTI T cells homed to the tumor site similarly at day 3 (Fig, 6H). At day 10, the control cells persisted whereas the Arhgef1-deficient cells disappeared (Fig. 6H). Notably, Arhgef1 was not required for OTI T cell accumulation in subcutaneously growing MC38 tumors; rather, the Arhgef1-deficient cells showed an advantage over WT cells (Fig. 6I). Taken together, these findings provide evidence that Gα13 function in CD8^+^ IEL contributes to promoting intestinal barrier integrity and anti-tumor immunity (Fig. S6C).

## DISCUSSION

Our findings establish a critical role for Gα13 and its downstream effector Arhgef1 in the homeostasis of intestinal CD8 T cells, including developmentally programmed CD8αα γδT cells and αβT cells and antigen-induced CD8αβ Trm cells. G13-pathway null cells show deficiencies in key gene expression programs and they undergo increased apoptosis. Imaging analysis shows Gα13 is needed to support IEL migration and positioning. We propose that Gα13 signaling supports intestinal CD8αα and CD8αβ T cell migration behaviors that are necessary for receipt of differentiation or survival signals, including IL15 for γδT cells and TGFβ for CD8αβ T cells. We identify GPR132 as a Gα13-coupled receptor that promotes IEL migration and contributes to CD8αβ IEL homeostasis. The strong intestinal T cell defect in Gα13-pathway deficiency is associated with exacerbated DSS-induced colitis and defective anti-tumor immunity. Based on the magnitude and breadth of the intestinal CD8 T cell loss caused by G13-pathway deficiency, and its selectivity to the intestine, targeting this pathway may have therapeutic potential for treatment of T cell driven intestinal inflammatory diseases.

Studies in IL15 and IL15Rα deficient mice established that IL15 is critical for γδT IEL and contributes to maintaining CD8αα αβT IEL.^49–52^ Our finding that the γδT IEL remaining in Gα13-deficient mice have reduced IL15 pathway activity suggests Gα13 supports migration behaviors needed for accessing IL15. Spatial gene expression analysis revealed IL15 is more abundant near the tips of villi than the crypts.^20^ The few γδT IEL remaining in Gα13^ΔVav^ mice were near the crypts, perhaps indicating poor access to IL15^+^ tips. Since IL15 promotes γδT IEL motility in vivo^39^, reduced access of Gα13-deficient γδT cells to IL15 may lead to a negative feedback loop and amplified cell loss. The inefficient movement of antigen-responding CD8αβ OTI T cells to the tips suggests a shared requirement for Gα13 in this migration behavior. Recent work established that IL7 cooperates with IL15 to maintain intestinal CD8αβ Trm IEL.^11^ IL7 is expressed by intestinal lymphatics and heterogeneously by intestinal epithelial cells, and spatial transcriptomics indicates greatest abundance mid-way along the crypt-villus axis.^20,53–55^ It is possible that limited access to both IL7 and IL15 contributes to poor survival of all Gα13-deficient CD8 IEL types. Reduced IL7 and IL15 signaling may contribute to reduced Bcl2 expression^11,56,57^ and thus increased apoptosis of G13-pathway deficient cells, a notion supported by our IL15–IL15Rα treatment study in the OTI *Lm*-OVA model.

TGFβR signaling in CD8αβ T cells has a well-established role in the maturation and maintenance of Trm IEL. In studies with LCMV infection and LCMV-specific P14 TCR transgenic T cells and OTI T cells in the *Lm*-OVA model, Tgfbr2^-/-^ cells arrive to the intestine in normal numbers one week after infection but then rapidly decay.^17–19^ We find the behavior of Arhgef1^-/-^ OTI cells in the oral *Lm*-Ova model has similar kinetics. In accord with a shared pathway, Arhgef1^-/-^ OTI cells in the intestine at day 8 have a reduced TGFβ gene expression signature and they can be partially rescued by TGFβ over-expression. The reduced TL binding by Gα13-deficient CD8αβ IEL, indicating reduced CD8αα expression, might also be a consequence of reduced TGFβ signaling.^16^ The sources of TGFβ needed for intestinal CD8αβ T cells are not clear though many cell types express one or more of the TGFβR ligands (TGFβ1, TGFβ2 and TGFβ3).^20^ A critical requirement for TGFβ to signal is that it must be activated from a latent state. TGFβ activators include integrins αvβ6 and αvβ8.^43^ Notably, αvβ6 is more abundant at the tips of intestinal villi than in the crypts (^58^ and this report). Moreover, spatial gene expression analysis showed higher amounts of TGFβ-induced gene transcripts in CD8 Trm cells nearer the tip.^20^ In mice lacking *Itgb6* (the gene encoding β6) and infected with LCMV, P14 cells reached the intestine in normal numbers at one week but were markedly reduced in the epithelium at day 42.^59^ While the exact sites of TGFβR engagement remain to be defined, we suggest that Gα13 and Arhgef1 are needed for CD8αβ T cells to position in a manner that enables adequate TGFβ exposure for their maturation and maintenance.

In our gene expression analysis of Gα13-deficient CD8αα αβT cells and Arhgef1-deficient CD8αβ Trm cells, the transcriptional regulators Nr4a2 and JunB were significantly less expressed. Using shRNA knockdown or conditional knockout, Nr4a2 and JunB were shown to be required for intestinal CD8 Trm and CD8αα IEL homeostasis.^60,61^ Gα13-pathway mediated induction of Nr4a2 and JunB may therefore contribute to IEL homeostasis.

The nuclear hormone receptor AHR is important for γδT and CD8αα αβT IEL homeostasis and contributes to CD8αβ Trm formation.^1,62,63^ We did not observe expression of several classical AHR targets^64^ in IEL such as *Cyp1a1*, *Cyp1b1*, or *Cyp1a2*, while *Ahrr* and *Tiparp* expression was not affected. However, several genes identified as AHR targets in in vitro generated CD8 Trm-like cells^63^, including *Itgae*, *S1pr5*, *Klrg1*, and *Eomes*, were differentially expressed in Arhgef1-deficient OTI d8 IEL, indicating a possible defect in AHR function in the absence of Gα13-pathway signaling. This might occur due to mispositioning and reduced access to AHR ligands or defects in AHR cooperating factors. One study reported Gα13 interacts with AHR-interacting protein (AIP) to promote AHR degradation.^65^ However, our data do not provide evidence for increased AHR pathway activity in the absence of Gα13, making it unlikely that Gα13 acts in IEL by directly destabilizing AHR.

The endogenous ligands acting on GPR132 are not well defined. While LPC can promote GPR132^+^ cell migration (^28–31^ and this report), it will be necessary to establish means to prevent LPC production to determine the role it plays in IEL homeostasis. Other ligands reported for GPR132 include lactate, N-palmitoylglycine, 9(S)-hydroxy-10,12-octadecadienoic acid, epoxyeicosatrienoic acid, oxylipins, and the microbiome derived molecule commendamide.^66–70^ It will be of interest to test if the contribution of GPR132 to IEL homeostasis depends on the microbiome. In accord with a role in IEL function, GPR132-deficiency exacerbated DSS-induced colitis.^71^

As well as promigratory actions in response to GPR132 ligands, Gα13 may guide cell migration in the villi by sensing cues that engage migration inhibitory responses. In addition to LPI ^21^, we found that LPA mediated IEL migration-inhibition in transwell assays. Although we did not detect an effect on IEL homeostasis of CRISPR-mediated Lpar5 or Lpar6 targeting, we do not exclude redundant actions of these receptors. Future studies in combined Gα13-coupled receptor deficient mice will be needed to address possible receptor redundancy.

How Gα13 supports the basal motility of IEL is an important question. We previously observed Gα13-dependent basal motility in germinal center B cells.^72,73^ It is possible that IEL (or other cells in the IEL preparation) are producing ligands for Gα13-coupled GPCRs that act in an autocrine or paracrine fashion to promote the basal motility. As well as its role downstream of Gα13-dominantly coupled GPCRs, Gα13 can be activated by Gαi-dominantly coupled GPCRs such as CXCR4.^74^ In this context, dual activation of Gαi and Gα13 is thought to reinforce ‘frontness’ and ‘backness’ (via Rac and Rho), respectively, and this in turn augments motility.^75,76^ It is possible that Gα13 in IEL generates cell polarity even in the absence of external stimuli. Notably, isolated CD8αα IEL have an amoeboid morphology compared to the round shape of isolated LN T cells^77^ consistent with an intrinsically motile state.

Our findings favor a model where the action of Gα13 in cell migration and positioning is key to its role in IEL. However, we do not exclude other possible mechanisms of Gα13 action. These could include signaling for gene induction^25,78^, supporting integrin activity needed for adhesive events^79,80^, influencing TCR signaling^81^, or limiting pAKT and mTOR activity^82,83^ to enable the cells to be maintained in a partially activated (poised) state ^60,84^. Reduced CD8αα expression and/or reduced access to TL might contribute to exaggerated TCR signaling in intestinal CD8αβ T cells and thereby lead to increased cell death.^32^ Some studies have noted interactions between Gα13 and Itk, a kinase that functions downstream of the TCR and has an important role in IEL homeostasis.^85,86^ However, Itk inhibition reduced α4β7 and CCR9 induction under gut tropic culture conditions^86^, a phenotype not observed for Gα13-deficiency, making it unlikely that Gα13 and Itk are acting in the same pathway in IEL.

γδT cell deficiency is associated with more severe DSS-induced colitis.^87–89^ Mice lacking αβT cells were also found to suffer more severe DSS-induced colitis.^88^ The exacerbated colitis observed in Gα13^ΔVav^ mice and Gα13^ΔCD4^ mice is in accord with these findings. γδT cells contribute to tissue repair by producing KGF1.^87,90^ New findings indicate that a subset of regulatory CD8 αβT IEL protects from colitis by suppressing macrophage activity.^7^ The constellation of mechanisms by which Gα13-dependent CD8 IEL protect from colitis requires further investigation.

Intrarectal implantation of the MC38 colon tumor line (derived from a C57BL/6 grade III adenocarcinoma^91^) provides a model for colorectal cancer and is associated with T, B and myeloid cell recruitment.^46–48^ Our findings indicate Gα13 pathway activity promotes CD8 T cell accumulation in colonic MC38 tumors. Importantly, Gα13 pathway deficiency had the opposite effect for subcutaneously growing MC38 tumors. The latter data are in accord with findings for Arhgef1-deficient CD8 T cells in a lung tumor model.^92^ Moreover, a recent genome wide CRISPR screen identified Gα13 as a negative regulator of human T cell accumulation in skin tumors in immunodeficient mice, an activity that may contribute to immune exclusion.^93^ We suggest that the dominant influence of Gα13 on tumor infiltrating T cell responses will differ by tissue site, with the essential role of Gα13 in intestinal IEL maturation and homeostasis playing a positive role in responses to intestinal tumors.

Using LCMV infection to broadly induce Trm cells, we found the Trm cell dependence on Gα13 was selective for the intestine, though we do not exclude roles in tissues we did not assess. The basis for this selectivity will require future exploration but it may in part reflect specialized migration requirements for intestinal Trm cells to access supportive niches. Beyond Trm cells, prior work on Gna13f/f CD4Cre mice identified a ∼2-fold deficiency of Tfh cells in response to immunization, LCMV infection or allergen exposure.^94^ Importantly, the impact of Gα13-deficiency on intestinal CD8 T cells was substantially more marked than the findings for Tfh cells. The selective dependence of intestinal CD8 T cells on Gα13 signaling makes this pathway of particular interest for therapeutic targeting in the context of T cell-driven inflammatory intestinal diseases.

### Limitations of Study

The very few IEL remaining in Gα13^ΔVav^ mice make it possible that some of the gene expression changes identified in the scRNAseq analysis are due to effects of niche emptying, Gα13-deficeincy in other cell types, or an altered microbiome. This caveat is less likely to apply to the RNAseq data collected from Gα13^ΔCD4^ and transferred Arhgef1^-/-^ OT1 cells. The studies with retroviral Bcl2 and activated TGFβ over-expression and with IL15-complex treatment support the model of Gα13 being required for access to trophic factors but as these are gain-of-function experiments, alternative explanations cannot be excluded. Our GPR132 findings are in accord with Gα13 being required for GPCR function, but it remains possible that some actions of Gα13 are independent of GPCR coupling. The observation of increased intestinal leakiness and reduced control of intestinal *C. rodentium* was only tested in Gα13^ΔVav^ mice, leaving open the possibility that Gα13 actions in cell types in addition to T cells contribute to these phenotypes.

## Resource availability

### Lead contact

Further information and requests for resources and reagents should be directed to and will be fulfilled by the Lead Contact, Jason Cyster.

### Materials availability

This study did not generate new, unique reagents.

### Data and code availability

RNAseq data have been deposited at GEO: GSE321684

## Acknowledgements

We thank Dario Vignali (University of Pittsburgh, PA) for E8iCreERT2 mice. We thank Molly Bassette from Matthew Krummel’s laboratory for the MC38-OVA cell line. We thank Vinh Nguyen and the UCSF Parnassus Flow Cytometry CoLab for help with cell sorting. We thank the UCSF genomics CoLab for help with RNAseq. ZME was supported by T32AI007334, T32AR079068, and is supported by a Cancer Research Institute Irvington postdoctoral fellowship. JGC is an Investigator of the Howard Hughes Medical Institute. This work was supported in part by NIH grant R01 AI040573 and NHLBI R35HL161241 and a Kenneth Rainin Foundation Health Innovator Award.

## Author contributions

Conceptualization: Z.M.E. and J.G.C.; methodology: Z.M.E. and J.G.C.; writing: Z.M.E. and J.G.C.; bioinformatic analysis: A.R., K.K., F.P.; formal analysis: Z.M.E., A.R., K.K., F.P; investigation Z.M.E., A.R., L.Q., N.J., A.K., W.L., H.T., J.A., Y.X., D.L.; resources: L.Y., M.R.L.; visualization: Z.M.E., A.R.; supervision: J.G.C.; funding acquisition J.G.C..

## STAR Methods

### 1. EXPERIMENTAL MODEL AND STUDY PARTICIPANT DETAILS

#### Animals

7-12 week old littermate mice of both male and female sex were used for experiments, and co-housed in specific pathogen-free conditions, at University of California San Francisco Laboratory of Animal Research Center. C57BL/6J, B6.SJL-*Ptprc^a^ Pepc^b^*/BoyJ, B6.Cg-*Commd10^Tg(Vav^*^1^*^-icre)A2Kio^*/J, Tg(Cd4-cre)1Cwi/BfluJ, B6.129S-*Tcrd^tm^*^1.1^*^(cre/ERT^*^2^*^)Zhu^*/J, B6.129P2-*Tcrb^tm1Mom^ Tcrd^tm1Mom^*/J, B6(C)-*Gt(ROSA)26Sor^em1.(CAG-cas9*,-EGFP)Rsky^*/J, B6.Cg-*Gt(ROSA)26Sor^tm^*^6^*^(CAG-ZsGreen^*^1^*^)Hze^*/J (Ai6), B6.Cg-*Gt(ROSA)26Sor^tm^*^14^*^(CAG-tdTomato)Hze^*/J (AI14), C57BL/6-Tg(TcraTcrb)1100Mjb/J were obtained from the Jackson Laboratory. Gna13fl/fl^95^ and Arhgef1-/- mice^80^ on a C57BL6/J background were from an internal colony. GPR132 deficient mice^45^, B6.129X1(C)-*Gpr132^tm1Witt^*/J, were a gift from Li Yang at East Carolina University and were originally generated at UCLA in the laboratory of Owen Witte. E8icreERT2 mice^36^ were a gift from Dario Vignali at University of Pittsburgh. Mice were maintained on 5058 -PicoLab Mouse Diet 20. For BM chimeras, mice were lethally irradiated by 450 rad x-ray irradiation in two-doses 3 hours apart and transferred retroorbital with at least 5×10^6 BM cells from donors of the indicated genotype. Mice were analyzed 8 weeks after reconstitution. All animal procedures were approved by the UCSF Institutional Animal Use and Care Committee.

#### Cell lines

Platinum-E (Plat-E) are an ecotropic retroviral packaging cell line derived from female human embryonic kidney 293T cells and stably express viral structural proteins (gag, pol, and env). They were used to generate MSCV or pTR retrovirus for overexpression in mouse T cells or BM. Cells were grown at 37°C, and 5% CO2 in DMEM supplemented with 10% fetal bovine serum, 2mM glutamine, 1% penicillin-streptomycin, 10mM HEPES, and 1μg/ml puromycin and 10μg/ml blasticidin. Cells were cultured to 80% confluency and then split 1:10. MC38 expressing OVA is an adenocarcinoma from female C57BL/6 mouse and was a gift from Matthew Krummel, UCSF. Cells were grown at 37°C, and 5% CO2 in DMEM supplemented with 10% fetal bovine serum, 2mM glutamine, 1% penicillin-streptomycin, and 10mM HEPES. Cells were cultured to 80% confluency and then split 1:10.

### 2. METHOD DETAILS

#### Tamoxifen treatments

Gα13fl mice were crossed to TCRδCreERT2 (Gα13^ΔTCRδ^) and E8iCreERT2 (Gα13^ΔCD8α^) and Ai6 or Ai14 reporters for inducible Ga13 knockout studies. We used homozygous TCRdCreERT2 mice to improve the deletion efficiency and heterozygous E8icreERT2. We first verified that our tamoxifen regimen of Cre– mice did not change IEL numbers (unpublished observations). To control for any Cre leakiness, we compared Cre expressing mice not treated with tamoxifen (Control) to Cre expressing mice (Gα13^ΔTCRδ^ or Gα13^ΔCD8a^) treated with tamoxifen. Tamoxifen (Sigma) was resuspended overnight at 20mg/ml in corn oil, shaking at 37°C in the dark, and 100ul was injected i.p. in the flank. Gα13ΔCD8α mice underwent 3 TAM treatments 24 hours apart for 6mg tamoxifen total, and Gα13^ΔTCRδ^ mice underwent 5 TAM treatments for 10mg total tamoxifen/ mouse. For two photon imaging studies, we utilized Gna13fl/+ TCRδCreERT2 Ai14 (Control) mice compared to Gna13fl/fl TCRδCreERT2 Ai6 (Gα13^ΔTCRδ^) mice.

#### MC38 cancer models

For MC38 orthotopic colorectal cancer studies, we utilized a published method of mechanical intestinal epithelial injury and rectal injection of tumor cells to facilitate colon tumor engraftment^48^. Prior to the procedure mice were fasted for 6 hours to ensure minimal colon content. Recipient mice, Gα13fl, Gα13^ΔCD4^, or TCRβδ^-/-^, were anesthetized with isoflurane and underwent rectal brushings with an interdental toothbrush dipped in EDTA that was prewarmed to 37°C to gently remove intestinal crypts. Crypts were examined under a light microscope to ensure similar brushings between mice. 2×10^6^ MC38-OVA cells were injected with a pipette tip intrarectally and a plug was created with ∼20ul of Vetbond Tissue Adhesive (3M) to prevent the cells leaking from the rectum. Mice were kept under 1% isofluorane for 10min with a warming pad, moved back to cage, and 4 hours later the plug was gently removed. Mice were weighed daily and weight loss relative to initial weight was calculated. Due to some inevitable inherent variability in brushing strength and tumor engraftment with this procedure, mice that did not have visible tumor engraftments one week after the procedure were removed from analysis. Tumor volumes were measured by an electronic caliper and calculated as V= 0.5*L*W^2^ where L= length and W= the width of the tumor. For subcutaneous MC38, 100ul were injected on each side containing 200,000 cells of MC38-OVA. Tumors were dissected, minced, and digested in 12ml of RPMI with 10% FBS, 1mg/ml Collagenase VIII (Sigma C2139), and 20ug/ml DNase1 while shaking at 37C for 30min. The remaining tissue was mashed over a strainer. The cells then underwent a 40% percoll density gradient centrifugation step to remove dead cells and debris.

#### Intravital microscopy of the small intestine

Mice were anesthetized with ketamine/xylazine (60/20 mg/kg, i.p.), and the abdominal area was shaved and prepared for surgery. Anesthesia was maintained with 1% isoflurane delivered in oxygen (1 L/min) through nosecone. Mice were positioned supine on a heating pad maintained at 37 °C. A small laparotomy was performed by incising the abdominal skin and underlying fascia to expose a short segment of small intestine. The mid-small intestine (jejunum or proximal ileum) was exteriorized and opened along the antimesenteric border using a cautery pen (Bovie high-temperature cautery pen, fine tip; ref. AA25) to expose the mucosal surface.

The tissue was gently repositioned with a saline-moistened cotton-tipped applicator such that the mucosal surface faced upward. A 3D-printed imaging window (prepared as previously described^96^) connected to a suction device was lowered onto the exposed tissue, and negative pressure (−20 mmHg) was applied to immobilize the intestinal segment against the coverslip during imaging.

Intravital imaging was performed on a Nikon A1R microscope equipped with a CFI75 Apochromat 25XC water-immersion objective and a high-resolution galvano scanner (UCSF Biological Imaging Development CoLab). Multiphoton excitation was provided by a Mai Tai DeepSee IR laser (950 nm). Emitted fluorescence was collected through 440/80-, 525/50-, and 600/50nm emission filters. Images were acquired at either 512 × 512 or 1,024 × 1,024 pixel resolution (when motion stability permitted). Time-lapse z-stacks spanning 9-12 µm were collected with 4-µm z-steps for 25-30 min. Raw image data were processed using the NIS.ai module in NIS-Elements (Nikon) to enhance signal-to-noise ratio.

Processed files were imported into Imaris 9.6.0 (Bitplane) for video rendering and cell tracking. IELs were identified and tracked using the built-in autoregressive spot-tracking algorithm. Tracks were filtered based on duration and total track length to exclude tracking artifacts. Automated tracking results were visually inspected and manually corrected when necessary. Mean track velocity was calculated directly from Imaris outputs and confinement ratio was defined as the ratio of track displacement to total track length (displacement/track length).

#### Bacterial infections

2×10^9^ CFUs of *Listeria monocytogenes*, strain *Lm*^InIA^-OVA, or 1×10^10^ CFU of kanamycin resistant *Citrobacter rodentium* DBS120 pler-lux, were orally inoculated into mice by gavage in 200ul of PBS after growing to an OD of 0.57 *(Lm*-OVA) or 0.72 (*Citrobacter*). *Lm*-OVA infections occurred the day after adoptive T cell transfers or 2-3 hours after when cells were sorted. Mice were analyzed 8 days after infection for effector response or 28 days after for memory T cell response. For quantifying CFUs of *Citrobacter* that translocated tissues were sterilely dissected and weighed, homogenized with a rotor in PBS, and then plated on MacConkey agar containing 50ug/ml of kanamycin to select for *Citrobacter rodentium*. 24 hours later CFUs were counted and quantified per g of tissue.

#### FITC-dextran permeability assay

FITC-dextran (Sigma) 4kd was dissolved in a stock of 100mg/ml in PBS and 15mg was gavaged to mice that were fasted for 4 hours. 4 hours later serum was collected, spun down, and quantified with a fluorescence spectrophotometer using a standard curve of FITC-dextran.

#### DSS colitis

2% DSS (MW ca 40000, Thermo Scientific) was administered ad libitum in the drinking water to Gα13^ΔVav^ and Gα13^ΔCD4^ and littermate control mice. Mice were kept on DSS water for five days, followed by regular water for three days and euthanized. Weight was recorded every day at the same time throughout the duration of the DSS protocol and data were graphed as % of initial weight.

#### Isolation of intestinal cells

To isolate IELs for downstream flow cytometry or sequencing analysis, 14 cm of the small intestine from the stomach was excised and left over segments were discarded. The entire colon was taken after the cecum to the rectum. The Peyers patches were removed, the intestine was opened longitudinally, cut into 1cm pieces, and added to Hanks’ Balanced Salt Solution media containing 10mM of HEPES buffer, 5mM of EDTA, and 1mM of DTT. The intestine pieces underwent two 20min shakes at 250RPM at 37C. The IEL cells were strained and then underwent a 40% percoll density gradient centrifugation step to remove dead cells and debris as previously described.^97^ LP cells underwent two further 20min shakes at 250RPM at 37C with RPMI, 10% FBS, 20ug/ml DNase1, and 100U/mg of Collagenase VIII (Sigma C2139). Following the two shakes the LP underwent a percoll density gradient spin similar to the IELs. A fraction of the IELs and LP cells were taken for flow cytometry staining. In some experiments mice were pre-treated 5 minutes before euthanasia with intravenous anti-CD45-PE (5ug) to label intravascular cells.

#### Flow cytometry

The cells were first stained with FC block (CD16/32) to block nonspecific binding, and then with Zombie NIR (Biolegend) for 15min at 4°C to exclude dead cells. Next the cells were stained for cell surface markers for 25min at 4°C. For AnnexV staining the cells were then stained with AnnexV-PE for 15min at 25°C in 1X binding Buffer (BD, 556454), washed and resuspended in the binding buffer and then the data were quickly acquired using Aurora Cytek. The antibodies used are indicated in Key resources table. The following antibodies and clones were purchased from Biolegend: CD45.2 BV421 (104), CD45.1 BV605 (A20), CD4 BV605 (GK1.5), CD44 PE-Cy7 (IM7), CD62L PE (MEL-14), Epcam PerCP-Cy5.5, AF488 (G8.8), CD3ε AF700 (145-2C11), CD8α APC-fire 750 (53-6.7), TCR Va2 APC (B20.1), CD103 PE-cy5 (2E7). The following antibodies and clones were purchased from BD: CD8β BV480, BV421 (H35-17.2), TCRβ BUV737 (H57-597), vβ5.1/5.2 TCR PE (MR9-4). The following antibodies and clones were purchased from Thermo Fisher: TCRγδ PE-cy7, AF647 (eBioGL3). Counting beads were added to determine absolute cell numbers in the original sample / amount of tissue taken. The cells were run on Aurora (Cytek) and data were analyzed using FlowJo software (Treestar). All cells were gated FSC SSC, singlets, and live. IELs were first gated: CD45^+^ CD3^+^. TCRγδ^+^ (TCRγδ IELs) or TCRβ^+^ then CD4^+^ CD8α^-^ (CD4 IELs). CD8α^+^ CD4^-^ then CD8β^+^ (CD8αβ IELs) or CD8β^−^ (CD8αα IELs).

#### Retroviral constructs and transductions

Bcl2 retroviral construct was made by inserting the murine open reading frame into the MSCV2.2 retroviral vector followed by an IRES and a GFP expression marker. The active TGFβ construct was generated by making a cysteine to serine substitution in human TGFβ1 at amino acid residue 33 ^42^ and cloned this into MSCV2.2 retroviral vector followed by an IRES and Thy1.1 reporter. For the G protein-coupled-receptor screen, single-guide RNA (sgRNA) sequences were cloned into the pTR-MSCV-IRES-Thy1.1 or pTR-MSCV-IRES-BFP vector.

SgRNA sequences were selected using Benchling’s CRISPR Guide tool and 2 guides were designed for each GPCR. To produce retroviruses, Platinum-E packing cells (Cell Biolabs) were grown to 75% confluency in a 10cm tissue culture dish and transfected with 10ug plasmid/25ul of Lipofectamine 2000 (Thermo Fisher) in OptiMEM following manufacturer’s protocol.

ViralBoost (Alstem) was added to the media 16 hours later and retrovirus containing supernatants were collected 48 and 72 hours after transfection, filtered through a 0.45um filter to remove debris, and concentrated 10 fold with Retro-X concentrator (Takara). For BM transductions, constitutive CAS9-GFP expressing donor mice were injected i.p. with 3 mg of 5-fluorouracil (Sigma), BM was collected after 4 days and cultured with complete DMEM supplemented with 20 ng/ml of interleukin-3 (IL-3), 50 ng/ml of IL-6, and 100 ng/ml of stem cell factor. BM cells were spin-infected (1000*g*, 2 hours, 32°C) at day 1 and day 2 and injected into irradiated recipients after the second transduction. For T cell transduction, CD8αβ T cells were activated with αCD3/CD28 T cell activator Dynabeads (Gibco) and 60U human IL-2 (NCI) in complete RPMI for 24 hours. The T cells underwent 2 spinfections (1000*g*, 2 hours, 32°C) with retrovirus 24 and 48 hours later on 20ug/ml Retronectin (Takara) coated 6 well plates.

#### RNP mediated gene targeting

CRISPR editing was performed via electroporation of Cas9/sgRNA ribonucleoprotein (RNP) complexes. Gna13-targeting sgRNAs were designed using CHOPCHOP to select the two highest ranked guides (GCAACGTGATCAAAGGTGCG and GAATTCCCGGCGCCGATCGT). As a negative control, an sgRNA targeting a sequence absent in both the human and murine genomes (GCGAGGTATTCGGCTCCGCG) was used. Lyophilized sgRNAs (Integrated DNA Technologies) were individually resuspended at 80 uM using 0.2 micron-filtered nuclease-free water. For each electroporation reaction, 1.25 uL of each sgRNA solution (or 2.5 uL of negative control sgRNA) were combined followed by the addition of 2.5 uL 40 uM Cas9 (UC Berkeley QB3 MacroLab) while slowly stirring with the pipette tip. The resultant 5 uL solution was incubated at 37°C for 20 minutes to induce RNP assembly, followed by a stable incubation at room temperature lasting no more than forty minutes. Cells were electroporated using the Lonza P3 Primary Cell 96-well Nucleofector Kit. Approximately 20-24 hours after activation, cells were washed with PBS and resuspended in P3 electroporation buffer at a concentration of 500,000 cells per 20 uL. For each reaction, 20 uL of cells were added to a 96-well Nucleocuvette Plate well. Each well was preloaded with the assembled RNP complex (5 uL) and a homology-directed repair template (1 uL of 100 uM stock solution; TTAGCTCTGTTTACGTCCCAGCGGGCATGAGAGTAACAAGAGGGTGTGGTAATATTACGGTACCGAGCACTATCGATACAATATGTGTCATACGGACACG) with no homology to the mouse genome to improve editing efficiency. Immediately after electroporation (Lonza 4D-Nucleofector Core Unit) with the pulse code EO-148, 80 uL of prewarmed complete RPMI was added to each well, followed by a 15-minute rescue incubation at 37C. Cells were subsequently transferred to 24-well plates and plated in 50 U/mL hIL-2 (NCI). After 3 days, a media swap was performed to complete RPMI containing 5 ng/mL recombinant murine IL-7 (Peprotech). After 1 day, these rested cells were washed with PBS and transferred to recipients.

#### Adoptive transfers

OTI T cells were enriched from the spleen, MLN, and iLN of Arhgef1^+/-^ or Arhgef1^-/-^ OTI donor mice from using negative selection anti-biotin beads (STEMcell Technologies) after staining with the following biotinylated antibodies: αCD4, αCD11b, αCD11c, αCD49b, αF4/80, αTCRγδ, αTER119, αGR1, αI-A/I-E, αB220. Cells were counted on a hemocytometer, and 50,000 OTI T cells were transferred to CD45.1 mice retro-orbitally. For Bcl2 mixed overexpression, 50,000 GFP^+^ EV-GFP or BCL2-GFP transduced OTI cells were sorted and mixed with 50,000 GFP– cells and transferred to CD45.1 mice retro-orbitally. 1×10^6 mixed congenically marked (CD45.1/.2 or CD45.2/.2) Arhgef1^+/-^ or Arhgef1^-/-^ OTIs were transferred to TCRβδ^-/-^ mice three days after MC38 engraftment retro-orbitally for tumor studies.

#### IL15/IL15Rα treatment

For IL15 agonist experiments, Arhgef1^+/-^ and Arhgef1^-/-^ OTI cells were transferred to CD45.1 WT mice and subsequently infected with *LM-Ova*. 6 and 7 days after infection, mice were injected i.p with 10ug in 200ul saline of ALT803 (MedchemExpress) a stable complex of an IL15 activation mutant, IL-15N72D, and IL15Rαsushi domain/Fc fusion (referred to as IL15/IL15Rα complexes). On day 8 after infection mice were there euthanized and analyzed for AnnexV, BCL2, and IF.

#### Transwell migration assay

IELs were counted and resuspended in migration medium (0.5% fatty-acid free BSA, 10mM Hepes RPMI) at 4 × 10^6^cells / ml. Transwell filters (6 mm insert, 5 μm pore size, Corning) were placed on top of each well, and 100 μl containing 4 × 10^5^ cells were added to the transwell insert. No ligand (Nil), lysophosphatidylcholine (LPC) 30uM, lysophosphatidic acid (LPA) 10uM, or CCL25 (1ug/ml) were added to the media at the bottom of the transwell. The cells were allowed to migrate for 3 hrs at 37°C with 5% CO2 and the cells in the bottom well were stained and counted by flow cytometry.

#### Histology

8 cm of jejunum or whole colon were excised and opened longitudinally. Intestinal content was removed. Starting from the most distal end and with the luminal side facing upwards, tissues were rolled into Swiss rolls and pinned together. For H&E staining, tissues were fixed with 10% formalin (Sigma-Aldrich), transferred to 70% ethanol 24 hours later, embedded in paraffin, and cut at 5 μm thickness at Histowiz laboratory, NY. For immunofluorescence staining, tissues were fixed with 1.5% PFA in PBS (Electron Microscopy Sciences) for 4-6 hours, washed in PBS, and transferred to 30% sucrose overnight at 4°C to dehydrate the tissue. The tissues were embedded in OCT and 1µum sections were cut with a cryostat. To stain the slides, the tissue sections were first washed for 10min in PBS, then permeabilized and blocked with 0.1% Triton X-100 (Sigma-Aldrich) 0.1% BSA and 5% normal rat and hamster sera (Jackson Immuno

Research) in PBS for 1 hour at room temperature. Antibodies were added to the blocking solution at 1:200 for TCRγδ-AF647, 1:200 CD8β-BV421, 1:500 EpCAM-AF488, and CD45.2 AF594 and stained overnight at 4°C. The slides were then washed three times with permeabilization buffer for 30 min total, briefly rinsed in water, and mounted with Aqua-mount (Thermo Fisher). The slides were imaged with Leica STELLARIS confocal microscope.

#### IF Imaging Analysis

Tiled images of 7 µm-thick jejunal sections were acquired using a Zeiss Axio Observer Z1 inverted microscope equipped with a Zeiss AxioCam 506 mono camera and a Zeiss Plan-Apochromat 10×/0.45 objective lens. Image acquisition and stitching were performed using ZEISS ZEN 2 software. Stitched images were imported into QuPath v0.6.0. for downstream analysis. Automated cell detection was performed using the QuPath cell detection module with fixed parameters across all samples. Detection settings were optimized to exclude artifacts based on cell size and signal intensity thresholds. To quantify CD45.2^+^ OT1 T cell distribution along the crypt-villus axis, line annotations (arrows) were manually drawn from the top of the crypt to the tip of the villus. Only villi meeting predefined inclusion criteria were analyzed.

Specifically, villi were required to: (i) exceed 70 µm in length; (ii) exhibit preserved morphology without horizontal or vertical collapse; (iii) be fully intact (not truncated); (iv) demonstrate EpCAM signal intensity allowing accurate crypt detection; (v) lack significant bending; (vi) not be in close proximity to adjacent villi that could confound assignment; and (vii) contain positively detected cells by QuPath. Using a custom script, each detected cell was assigned a relative position along its nearest villus axis via vector projection.

Cells located more than 50 µm from the annotated axis were omitted from analysis. The projected position of each cell was normalized along the length of the villus axis (0-1) and categorized into three bins representing spatial compartments: Low (0-.333), Medium (.333-.666), and High (.666.-1) villus regions.

#### Cell sorting for RNAseq

For cell sorting, 4000 TCRβδ^+^ TCRαβ^−^, 4000 TCRαβ^+^ CD8α^+^ CD4^-^ and 4000 CD4^+^ IELs, were sorted with Aria Fusion (BD Biosciences) from 3 pooled Gα13fl or Gα13^ΔVav^ mice that were hash-tagged during the surface staining to distinguish genotypes using CD45/MHCI (Biolegend TotalSeq). Samples were loaded on 10x Chromium Controller (10X Genomics) immediately after sorting. For bulk sequencing of CD8αβ IELs 100 TCRαβ CD8αβ^+^ IELs were sorted into SmartSeq 1X Reaction mix with 3’ SMART-Seq CDS Primer II (TAKARA). Libraries were generated with SMART-seq v4 (TAKARA) at UCSF Genomics facility and sequenced on NovaSeq (Illumina).

#### RNAseq

For scRNAseq, samples were run on a 10X Chromium chip with 3′ v.2 chemistry (10X Genomics) as per the manufacturer’s instructions by the UCSF Institute for Human Genetics Sequencing Core. Transcripts captured in all the cells encapsulated with a bead were uniquely barcoded using a combination of a 16 bp 10x Barcode and a 10 bp unique molecular identifier (UMI). cDNA libraries were generated using the Chromium™ Single Cell 3’ Library & Gel Bead Kit v3 (10x Genomics) following the detailed protocol provided by the manufacturer. Libraries were sequenced with the NovaSeq 6000 platform (S1 Cartridge, Illumina) in 150 bp paired-end mode. The hashtag library was demultiplexed using CellRanger software (version 7.0.0) ^98^.

Aligned reads were used to quantify the expression level of mouse genes and generation of gene-barcode matrix. Subsequent data analysis was performed using Seurat R package (4.0.4).^99^ Quality control was performed, and viable cells were selected by excluding cells with features lower than 1500 and above 4000, as well as cells having more than 3.5% of mitochondrial transcripts and more then 15000 transcripts. Control and Gα13^ΔVav^-derived cells were demultiplexed with the HTODemux function integrated in Seurat with standard settings. All doublets were removed for the analysis. 2000 most variable genes used for the anchoring process were used for downstream analysis to calculate principal components, after log-normalization and scaling. Principle component analysis (PCA) was used for dimensionality reduction and to visualize a uniform manifold approximation and projection (UMAP) of the identified clusters. P-values comparing gene expression of clusters and samples were calculated with the FindMarker function in Seurat. Gene list was generated using an adjusted p-value cutoff <0.05 and expression in the indicated cluster cutoff >25. For GSEA of TGFβ and IL15 induced genes, log2 fold change (LFC) and adjusted p values (padj) were given as input to the fgsea() function from the fgsea R package (v.1.32.0), which implements a pre-ranked gene set enrichment analysis. The rankings of the genes were based on generated score calculated by LFC * -log10(padj).

Raw bulk RNA-seq data were processed using STAR ^100^ and RSEM ^101^ in default settings. In short, raw fastq files were mapped to mm10 genome, and TPM was calculated using RSEM. Then the data was loaded into and further analyzed by DESeq2.^102^ Differential expressed genes were called with padj < 0.05 and abs(log2Foldchange) >= 0.6 (approximately fold change of 1.5). Enrichr ^103^ was used to perform pathway analysis of up-regulated genes and down-regulated genes, respectively, and selected significant biological pathways padj < 0.05 were plotted from MsigDB hallmark, WikiPathways, Gene Ontology, and KEGG databases.

Xenium spatial transcriptomics data were obtained from the publicly available dataset generated by.^20^ Imaging files and transcript coordinate data from small intestinal tissue at day 8 post-LCMV infection were processed and plotted using the public analysis pipeline developed by the Goldrath Lab (available on GitHub).

## 3. QUANTIFICATION AND STATISTICAL ANALYSIS

Prism software (GraphPad 9.0.1) was used for statistical analyses. Data were first analyzed for normal distribution using D’Agostino and Pearson omnibus normality tests and subsequently analyzed with two-tailed unpaired t-tests when comparing two groups, one-way ANOVA with Tukey’s post-hoc test for multiple comparisons, Fishers exact test for association with categorial variables to determine statistical significance. Data in all figures were pooled from a minimum of two-independent experiments and represented as mean ± SEM when possible. The statistical test used, sample size, and *P* values are indicated in each figure legend. *P* values <0.05 were considered statistically significant. \*\*\*\**P*<0.0001, *** *P*<0.001, ** *P*<0.01, * *P*<0.05.

### 5. KEY RESOURCES TABLE

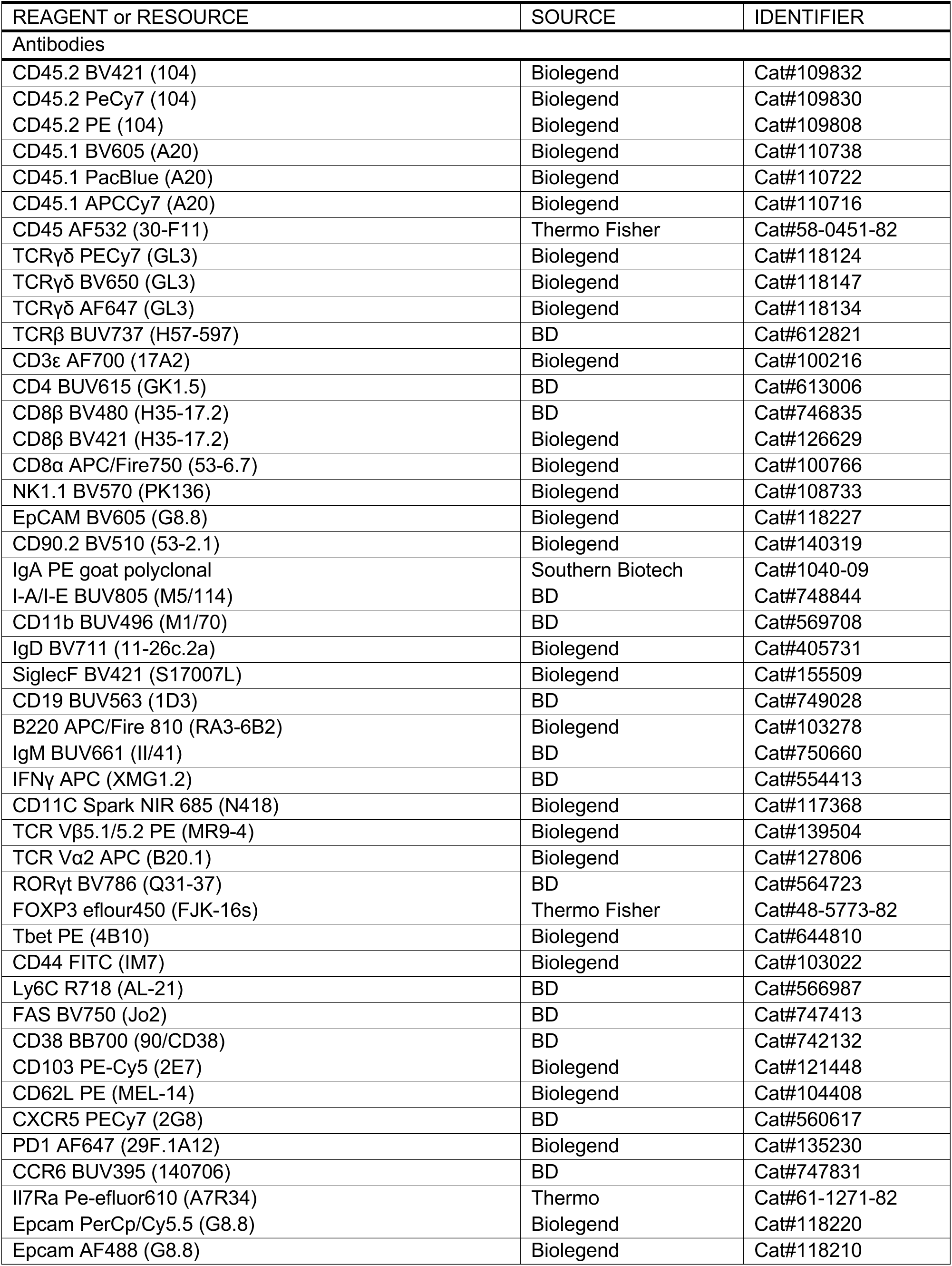
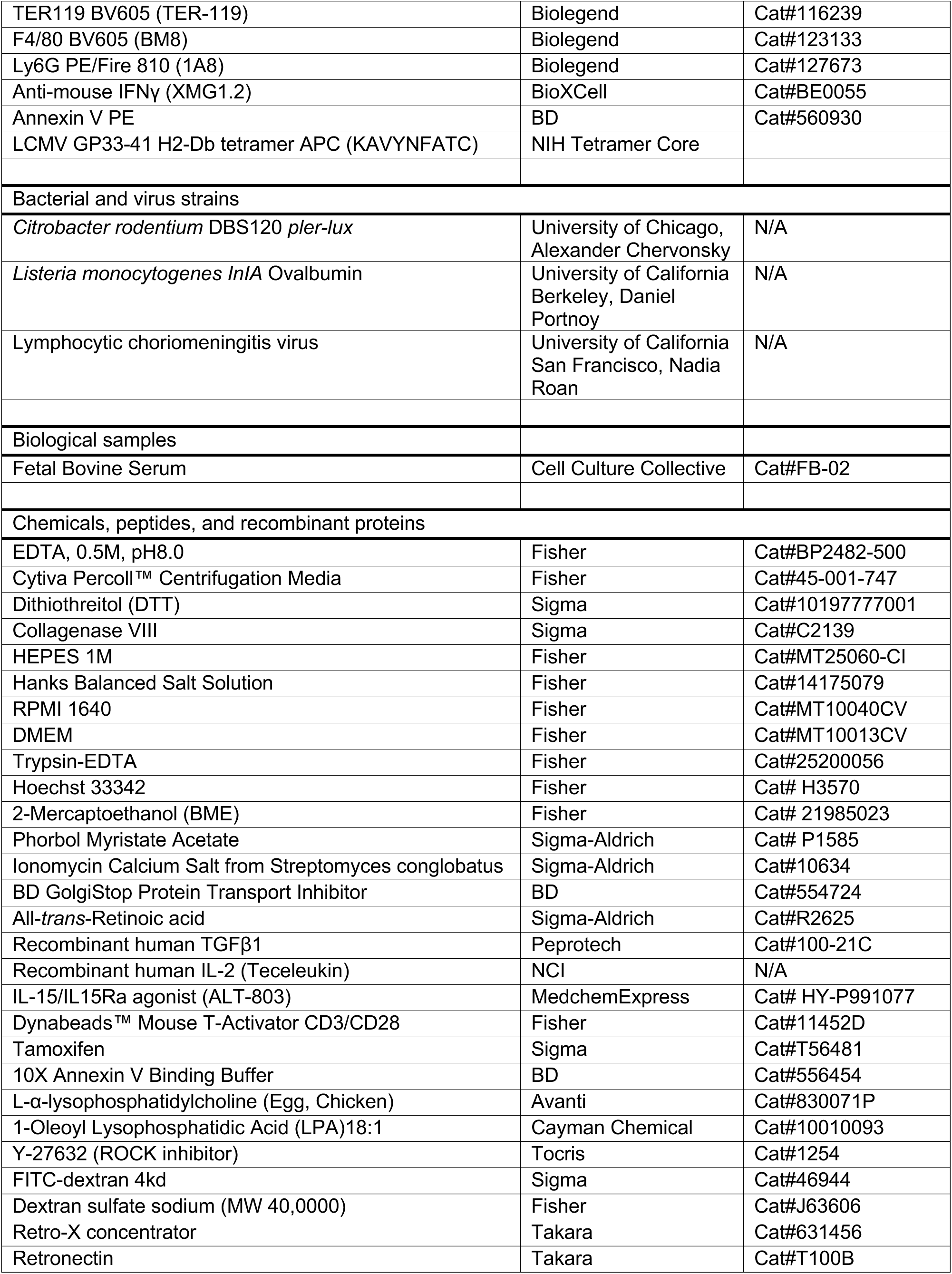
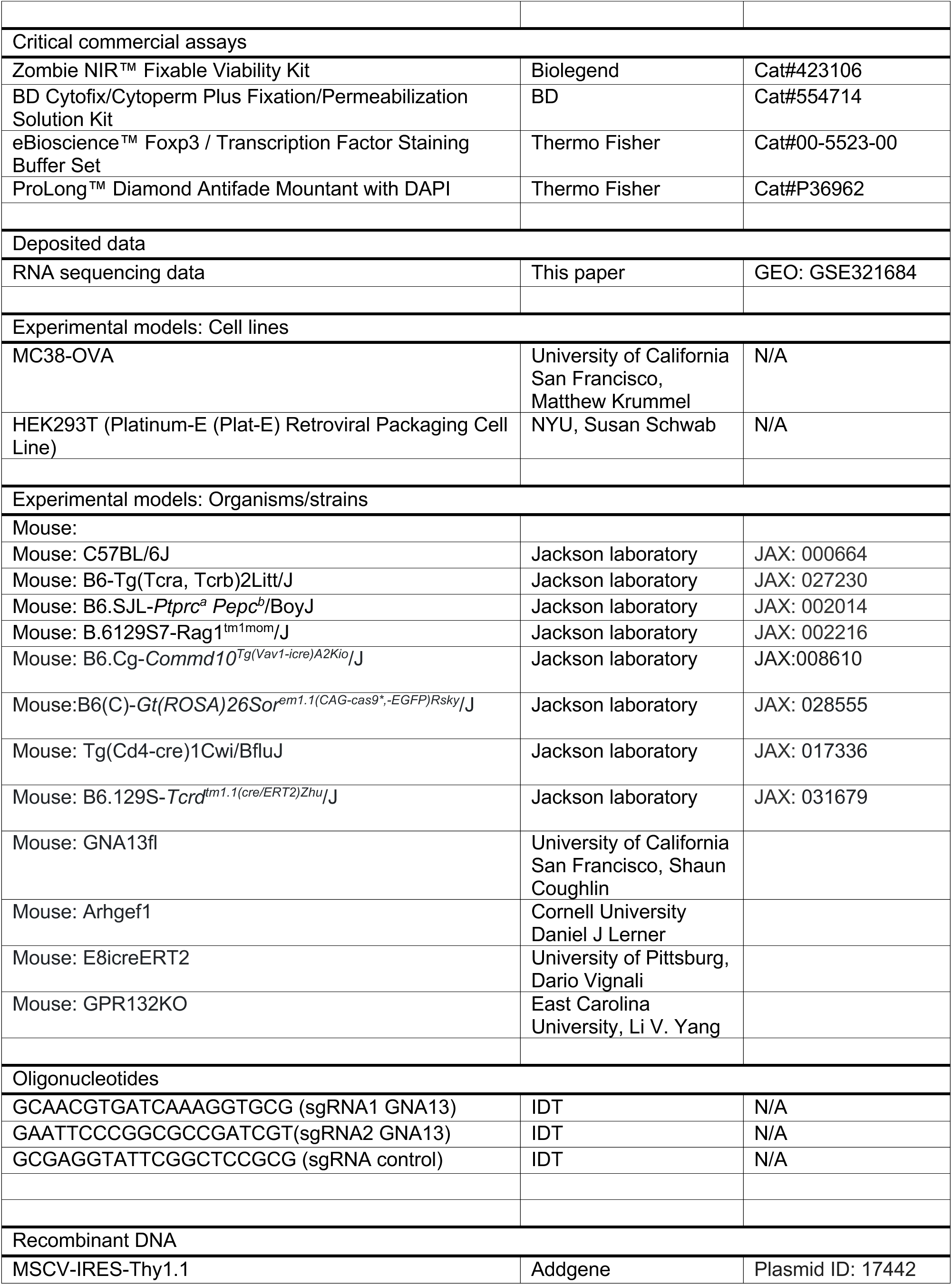
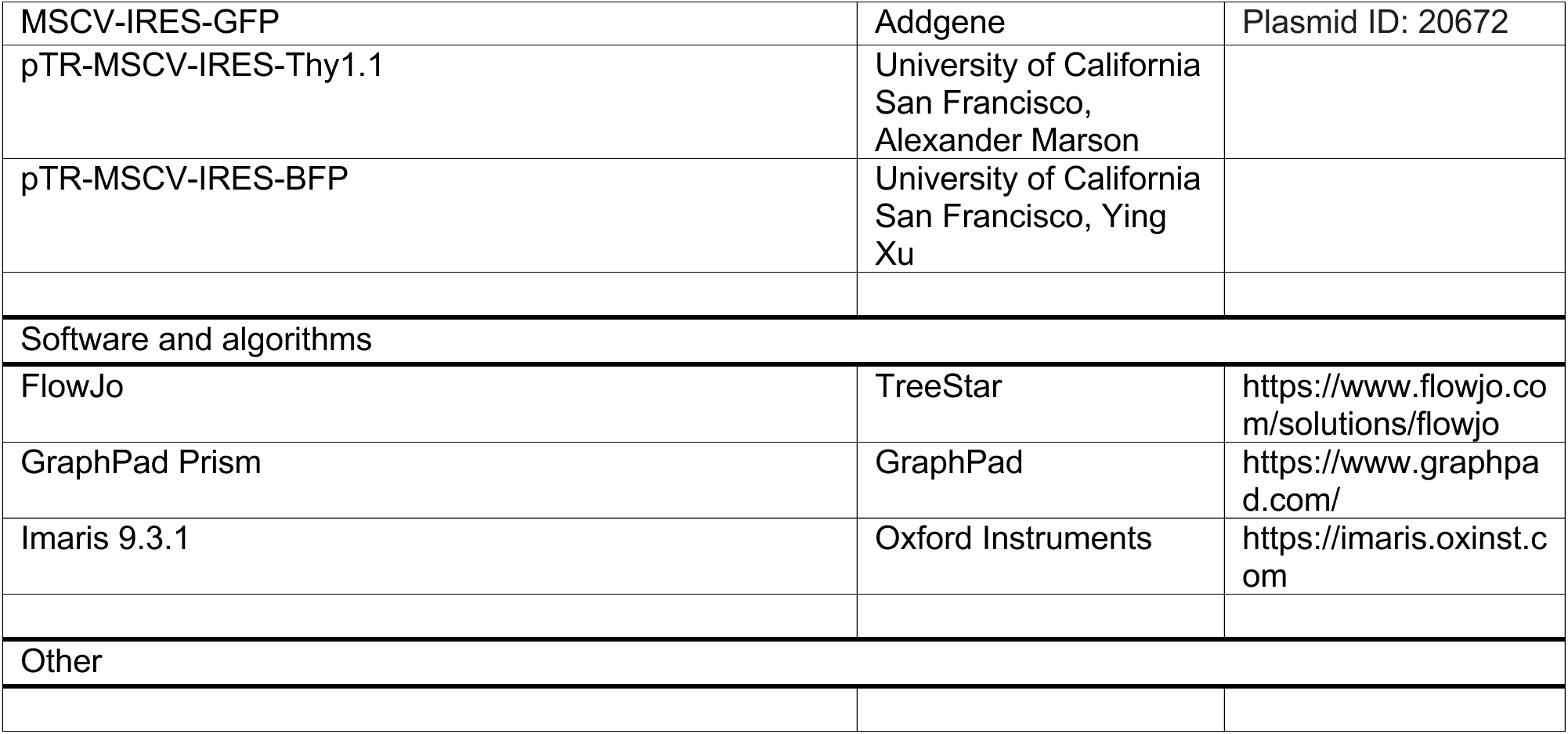

## Supplemental Information

Supplemental information includes 6 figures, 2 videos, and 1 table.

## Supplemental Movie Legends

**Movies S1. Real-time imaging of γδT IEL migration in small intestine of a Gna13f/f Ai6 TCRδCreERT2 mouse at various days after tamoxifen treatment.** The movie is a concatenated series of 20-30-min time-lapse sequence (duration indicated on bottom right of movie) of 9-12µm *z*-projection images. ZsGreen^+^ IEL are in green and nuclei in blue (Hoechst). The epithelium surrounding the villi in the projection image can be identified by the uniform lines of labeled nuclei. The nuclei within the interior of the villi help identify the lamina propria. Time after tamoxifen treatment: days 3, 7, 10, 14.

**Movies S2. Real-time imaging of γδT IEL migration in small intestine of a Gna13f/f Ai6 TCRδCreERT2 / WT Ai14 TCRdCreERT2 mixed BM chimeras at day 3 and 14 after tamoxifen treatment.** The movie is a concatenated series of 15-30-min time-lapse sequence of 9-12µm *z*-projection images with multiple fields of view (FOV). ZsGreen^+^ Gα13^ΔTCRδ^ IEL are in green, TdTomato^+^ WT **γδ**T IEL are in pink and nuclei in blue (Hoechst). The white lines show cell tracks. In some cases, the cell movement was too little to generate a visible track. TAM day 14 FOV 3 shows a zoomed in view highlighting an example of a Gα13^ΔTCRδ^ IEL that stopped moving during the imaging period.

Table 1: Antibodies used for Cytek analysis for figure 1

