## Supplemental figures for "GPCR-Gα13 signaling regulates survival of intestinal intraepithelial CD8 lymphocytes through migration to cytokine-rich niches"

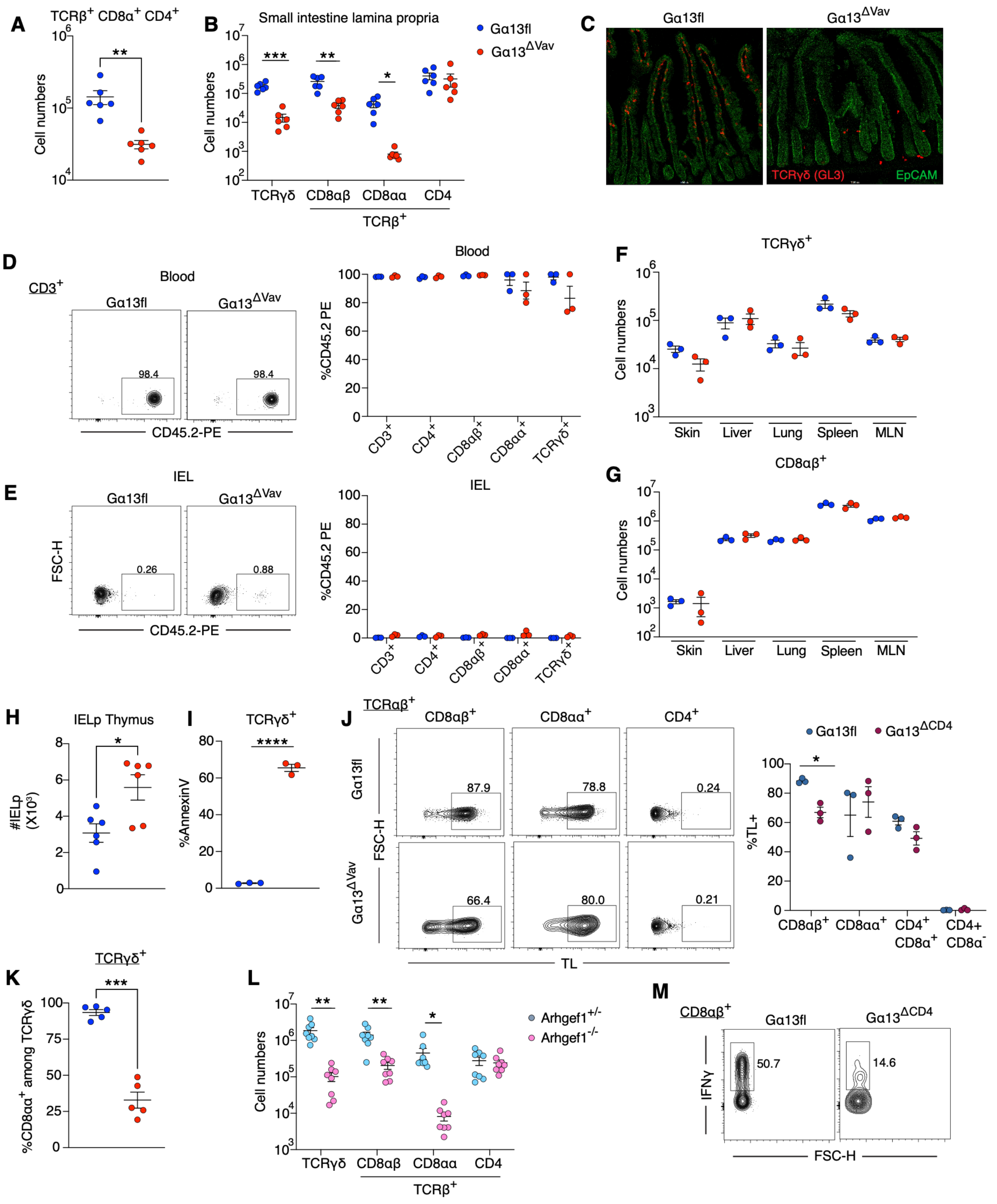


**Figure S1. Gα13 is required for intestinal IEL T cell homeostasis (related to Figure 1)**

A. Number of CD4 and CD8α double positive TCRαβ+ IELs in the small intestine of Gα13fl and Gα13$^{\Delta Vav}$ mice. (N= 6 mice/group).

B. Small intestine LP cell numbers of T cell subsets in Gα13fl and Gα13$^{\Delta Vav}$ mice. (N= 6 mice/group).

C. Representative IF image from the small intestine of Gα13fl and Gα13$^{\Delta Vav}$ mice stained for TCRγδ (red) and EpCAM (green).
D, E Intravascular labeling with CD45.2-PE and assessment of CD45.2-PE+ T cells in blood (D) or IEL (E) of Gα13fl and Gα13$^{\Delta Vav}$ mice. Left is representative FACS plot of CD45.2-PE staining in CD3$^+$ cells from blood and IEL. (N= 3 mice/group)
F, G. Numbers of TCRγδ$^+$ (C) or CD8αβ (D) T cells in the skin, liver, lung, spleen, and MLN of Gα13fl and Gα13$^{\Delta Vav}$ mice. (N= 3 mice/group).
H. Number of CD8αα IEL precursors in the thymus of Gα13fl and Gα13$^{\Delta Vav}$ mice gated: CD4- CD8α$^-$ TCRβ$^+$ CD5$^+$ CD122$^+$ H2kb$^+$ PD1$^+$ CD44$^{lo}$. (N= 6 mice/group).
I. Frequency of Annexin V$^+$ cells among TCRγδ$^+$ IEL. (N= 3 mice/group).
J. TL tetramer staining of TCRαβ$^+$ small intestinal IELs in Gα13fl and Gα13$^{\Delta CD4}$ mice. (Left) Representative FACS plot. (Right) Summary data of %TL$^+$ among CD8αβ$^+$, CD8αα$^+$ CD4$^-$, CD8α$^+$ CD4$^+$, and CD8α$^-$ CD4$^+$ cells. (N= 3 mice/group)
K. Frequency of CD8αα$^+$ among TCRγδ cells in Gα13fl and Gα13$^{\Delta Vav}$ mice. (N= 5 mice/group)
L. Total numbers of small intestinal IEL subsets in Arhgef1$^{+/-}$ and Arhgef1$^{-/-}$ mice. (N= 8 mice/group).
M. Representative FACS plot of IFNγ$^+$ CD8αβ IELs in Gα13fl and Gα13$^{\Delta Vav}$ mice.
All graphed data are pooled from two independent experiments and represented as mean ± SEM. ****$P$<0.0001 , *** $P$<0.001, ** $P$<0.01, * $P$<0.05, t-test.

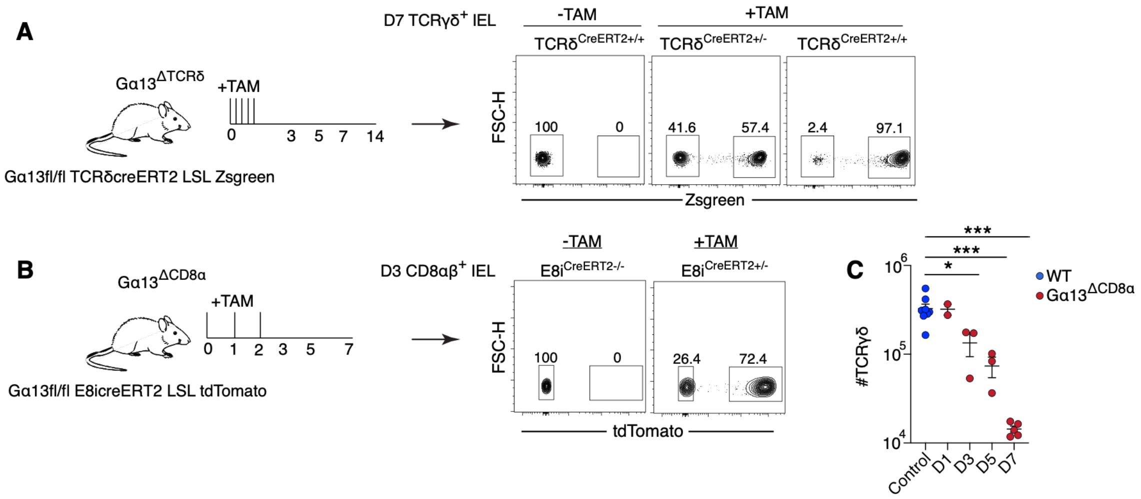


**Figure S2. Conditional Gα13 deletion from IEL leads to cell loss (related to Figure 2)**

A. Experimental scheme of tamoxifen (TAM) treatment in $G\alpha13^{\Delta TCR\delta}$ mice (left) and representative FACS plots of ZsGreen induction in TCRγδ IELs of homozygous and heterozygous $CreERT2^{+}$ mice 7 days after start of TAM treatments.
B. Experimental scheme of tamoxifen treatment in $G\alpha13^{\Delta CD8\alpha}$ mice (left) and representative FACS plots of tdTomato induction in CD8αβ IELs of heterozygous $CreERT2^{+}$ mice 3 days after start of TAM treatments.
C. Number of TCRγδ IELs in Ga13ΔCD8α mice, 1, 3, 5, and 7 days after start of TAM. (N= 2-8 mice/group).

Data in C pooled from two independent experiments and represented as mean ± SEM.

****$P$<0.0001 , *** $P$<0.001, ** $P$<0.01, * $P$<0.05, t-test.

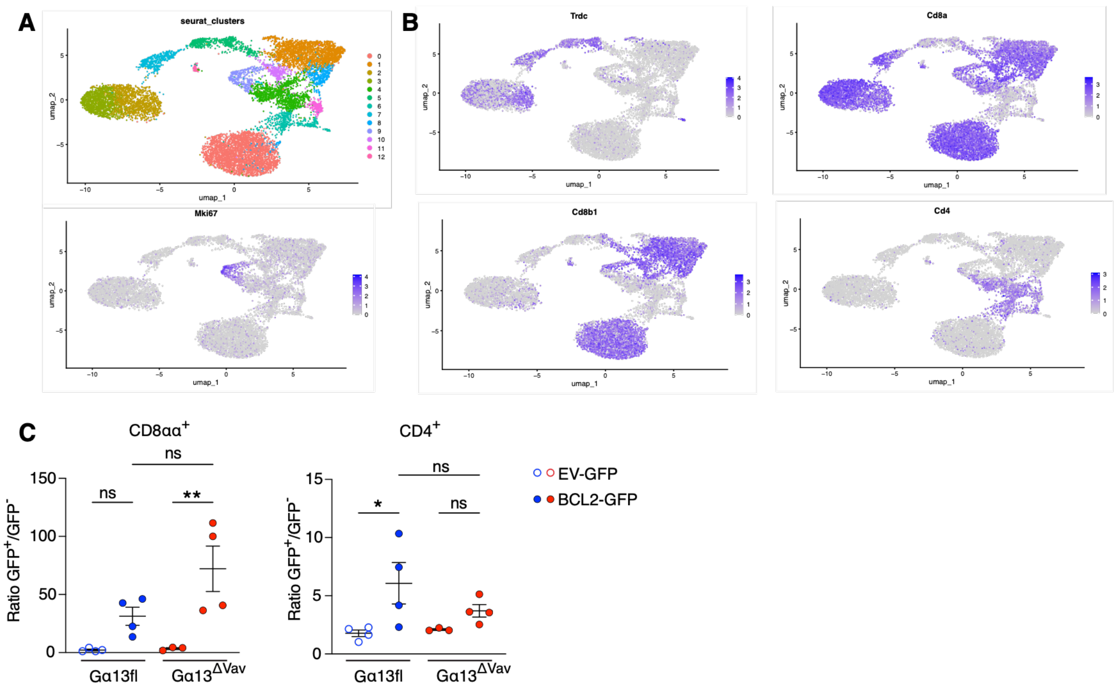


**Figure S3. Single cell RNA-seq analysis of Gα13-deficient intestinal T cells (related to Figure 3)**

A. uMAP of Seurat clusters from 10x RNAseq analysis of sorted IELs from Gα13fl and Gα13$^{\Delta Vav}$ mice.

B. uMAP of gene expression plots of *Tcrd*, *Cd8a*, *Cd8b1*, *Cd4* and *Mki67* from (A).

C. Retroviral BM chimeras with overexpression of EV-GFP (open circles) or Bcl2-GFP (filled circles) in Gα13fl (blue) and Gα13$^{\Delta Vav}$ (red) cells. Data are expressed as a ratio of GFP$^{+}$/ GFP$^{-}$ of each group of TCRαβ$^{+}$ CD8αα$^{+}$ (left) or CD4$^{+}$ (right) IELs. N = 3-4 mice/group and are representative of two independent experiments.

Data in C pooled from two independent experiments and represented as mean ± SEM. ** *P*<0.01, * *P*<0.05, ANOVA with Tukey multiple comparisons.

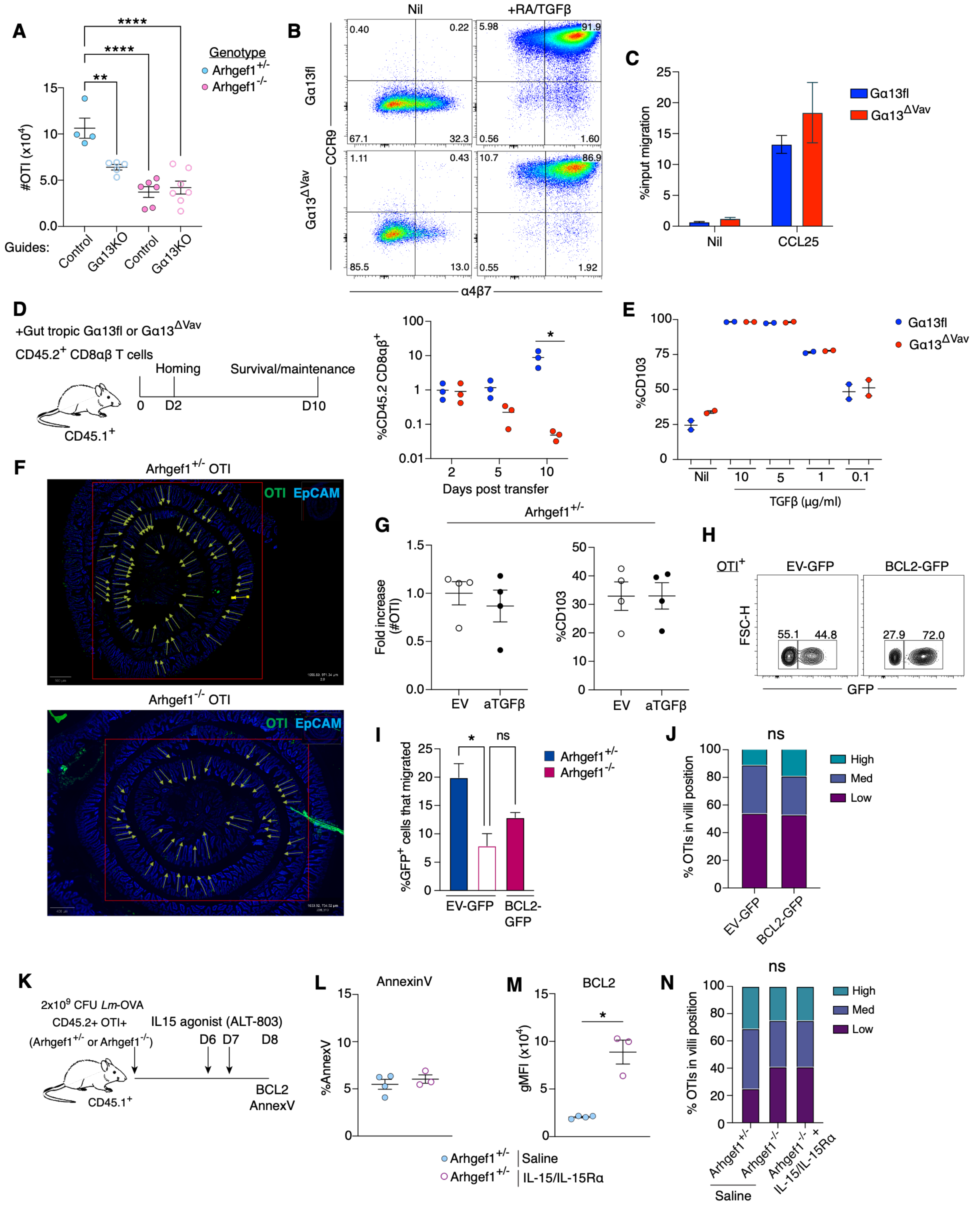

**Figure S4. Intact induction of gut tropism but poor intestinal survival of Gα13-deficient CD8αβ T cells (related to Figure 4)**

A. Number of OTI in small intestine IEL from CAS9 RNP knockout of non-targeting control or Gna13 guides in Arhgef1$^{+/-}$ or Arhgef1$^{-/-}$ OTI cells. (N=4-7 mice/group).
B. FACS plot of CCR9 and α4β7 levels in Ga13$^{\Delta Vav}$ CD8αβ T cells 5 days after activation in the absence (Nil) or presence of retinoic acid (RA) and TGFβ.
C. Frequency of CD8αβ IEL transwell migration to Nil or CCL25 of Ga13fl or Ga13ΔVav T cells 5 days after activation in gut tropic conditions.
D. Experimental scheme (left) of adoptive transfer of skewed gut tropic CD8αβ T cells generated as in (A) to CD45.1 WT recipients and analyzed 2, 5, and 10 days after transfer. Frequency of Gα13fl or Gα13$^{\Delta Vav}$ CD45.2 CD8αβ IELs recovered after transfer (right). (N=3 mice/group)
E. Frequency of CD103 upregulation in CD8αβ T cells stimulated with TGFβ in media from Gα13fl or Gα13$^{\Delta Vav}$ mice. (N=2 mice/group).
F. Example images used for quantitating transferred control or Arhgef1$^{-/-}$ OTI T cell distribution in small intestine of *Lm*-OVA infected recipients at day 8. Arrows indicate villi used for quantitation of CD45.2$^{+}$ OTI cells (green). Epithelium was detected with EpCAM (blue).
G. Retroviral overexpression of aTGFβ-Thy1.1 and control EV-Thy1.1 in Arhgef1$^{+/-}$ OTI T cells from Figure 4I. Left panels are fold increase in the absolute number of reporter+ cells, right graph shows frequency of CD103 in reporter+ OTI small intestinal IELs. (N=4 mice/group).
H. Representative FACS plot of IEL from mice that received Arhgef1$^{-/-}$ OTI cells transduced with control EV-GFP or Bcl2-GFP mixed equally with non-transduced cells, at day 11 post transfer to *Lm*-OVA infected hosts.
I. Transwell migration assay of Arhgef1$^{+/-}$ OTI IELs transduced with EV-GFP and Arhgef1$^{-/-}$ OTI transduced with EV-GFP or BCL2-GFP. (N=4 mice/group).
J. Frequency of Arhgef1$^{-/-}$ OTI transduced with EV-GFP or BCL2-GFP found in the low, med, and high position of the villi. (N=3 mice/group).
K. Schematic of IL15 agonist (ALT-803) treatment after Arhgef1$^{+/-}$ or Arhgef1$^{-/-}$ OTI transfer.
L. Geometric mean fluorescence intensity of BCL2 in Arhgef1$^{+/-}$ or Arhgef1$^{-/-}$ OTI cells from mice treated as in K. (N=3-4 mice/group).
M. Frequency of AnnexinV$^{+}$ Arhgef1$^{+/-}$ or Arhgef1$^{-/-}$ OTI cells from mice treated as in K. (N=3-4 mice/group).
N. Frequency of Arhgef1$^{+/-}$ or Arhgef1$^{-/-}$ OTI T cells found in the low, med, and high position of the villi from mice treated as in K. (N=3 mice/group).
All graphed data are pooled from two independent experiments and represented as mean ± SEM. ****$P$<0.0001 , *** $P$<0.001, ** $P$<0.01, * $P$<0.05, t-test, or ANOVA with Tukey multiple comparisons (A, I, M), or Fishers exact test (N).

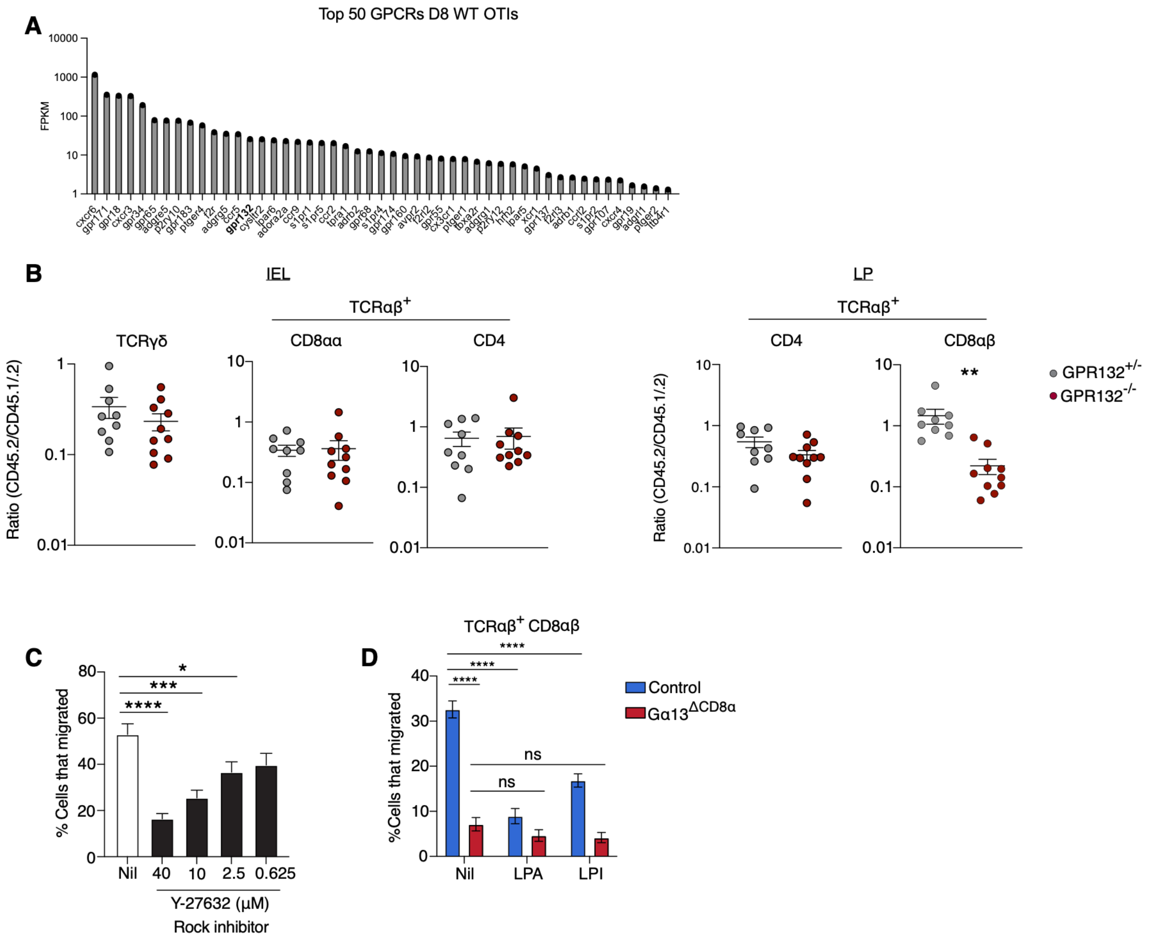


**Figure S5. GPCR expression in CD8αβ IEL, impact of GPR132-deficiency and migratory response to lysophospholipids (related to Figure 5)**

A. Top 50 GPCRs by FPKM expression from RNAseq of transferred WT OTI IELs 8 days post *Lm*-OVA infection.

B. Ratio of the indicated T cell subsets in the IEL and LP from BM chimeras made with $GPR132^{+/-}$ or $GPR132^{-/-}$ (CD45.2) BM mixed 1:1 with WT (CD45.1/.2) BM. (N=9-10 mice/group). Corresponds to mice shown in Fig. 5B.

C. Transwell migration assay of CD8αβ IEL incubated with no inhibitor (Nil) or Y-27632 (ROCK inhibitor) at the indicated concentrations. Migration was performed in the absence of chemoattractant. (N=3 mice/group).

D. Frequency of CD8αβ IEL transwell migration to Nil, LPA, or LPI, of control or inducible $G\alpha 13^{\Delta CD8\alpha}$ cells 5 days after TAM treatment. (N=4 mice/group).

Data in B, C, D are pooled from two independent experiments and represented as mean ± SEM. ****$P$<0.0001 , *** $P$<0.001, ** $P$<0.01, * $P$<0.05, t-test.

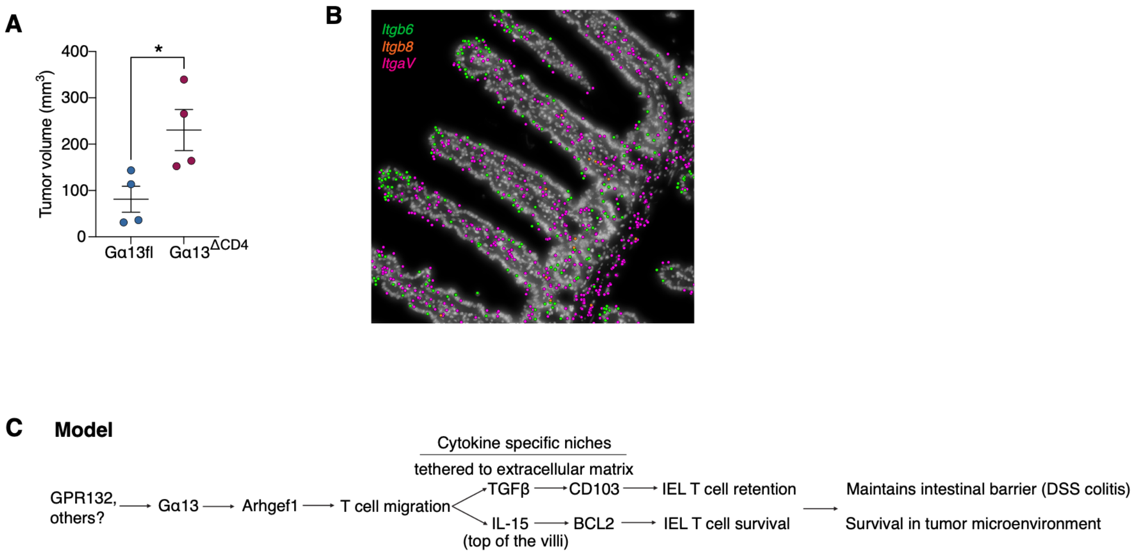


**Figure S6. Gα13 signaling in IEL T cell homeostasis (related to Figure 6)**

A. Tumor volume 7 days after implantation of colorectal MC38 in Gα13fl or Gα13$^{\Delta CD4}$ mice. (N=4 mice/group). Data are pooled from two independent experiments and represented as mean ± SEM. * $P$<0.05, t-test.

B. Distribution of transcripts for TGFβ-activating integrin chains *Itgav*, *Itgb6* and *Itgb8* in spatial RNAseq analysis of mouse small intestine at day 8 of LCMV infection, determined using data from [20]. Very few *Itgb8* transcripts were detected.

C. Model of GPCR-Gα13 signaling role in regulating IEL T cell homeostasis.
